# The *Chlamydia trachomatis* secreted effector CebN targets nucleoporins and Rae1 to antagonize STAT1 nuclear import

**DOI:** 10.1101/2024.04.25.587017

**Authors:** Brianna Steiert, Cherilyn A. Elwell, Xavier Tijerina, Madison A. Elliott, Yennifer Delgado, Jocelyn Ni, Robert Faris, Parker Smith, Shelby E. Andersen, Paige N. McCaslin, Julia Kevil-Yeager, Quinn Eldridge, Brian S. Imai, Justine V. Arrington, Peter M. Yau, Kathleen M. Mirrashidi, Jeffrey R. Johnson, Erik Verschueren, John Von Dollen, Gwendolyn M. Jang, Nevan J. Krogan, Mehdi Bouhaddou, Christine Suetterlin, Joanne N. Engel, Mary M. Weber

## Abstract

To usurp host defenses and establish a replicative niche, obligate intracellular pathogens are tasked with remodeling the host cell using a comparatively small repertoire of effector proteins. For *Chlamydia trachomatis* (*C.t*), discovery of secreted proteins and their host targets has been particularly challenging due to the bacterium’s historical genetic intractability. Using affinity purification-mass spectrometry, we defined host interaction partners for 21 secreted effector proteins, providing the first comprehensive type III secretion system (T3SS) effector- host interactome generated during infection. Among these, we show that the C-terminus of CebN (CT584) binds multiple nucleoporins and Rae1, host factors previously associated only with viral immune evasion. Remarkably, we shown that CebN localizes to the nuclear envelope not only in infected cells but also in uninfected bystander cells. Functionally, CebN is both necessary and sufficient to perturb STAT1 nuclear import following IFN-γ stimulation and its expression is critical for *C.t.* survival, as evidenced by reduced bacterial replication and smaller inclusions in cells infected with a CebN mutant. Together, these finds expand our understanding of chlamydia effector biology and highlight novel bacterial strategies for manipulating host defenses at the nuclear pore.

**SIGNIFICANCE:** *Chlamydia trachomatis* (*C.t.*) is a leading cause of sexually transmitted infections and blindness, yet the molecular mechanisms it uses to manipulate host defenses remain poorly defined. Unlike many pathogens, *C.t.* relies on a limited set of effectors to remodel the host cell and establish its niche. We identified host targets for 21 *C.t.* effector proteins. Focusing on CebN, we show that it binds nucleoporins and Rae1, host factors previously linked only to viral immune antagonism. CebN localizes to the nuclear envelope of both infected and bystander cells, and is critical for replication, inclusion development, and perturbation of STAT1 nuclear import following IFN-γ stimulation. These findings uncover a novel strategy by which *C.t.* manipulates nuclear pore function to evade host defenses and establish infection.

## INTRODUCTION

To counteract host defense mechanisms and establish a favorable replicative niche, intracellular pathogens are tasked with remodeling the host cell using secreted virulence factors, termed effector proteins. Identification of the host proteins and pathways targeted by secreted proteins during active infections has been exceedingly difficult for obligate intracellular pathogens owing to their genetic intractability^1^. Several obligate intracellular pathogens, including *Chlamydia trachomatis* (*C.t.*), are the etiological agents of important human diseases for which no vaccines exist^2^. *C.t.* is the leading cause of infectious blindness and is the most common bacterial sexually transmitted infection worldwide^3^. Untreated infection can result in severe complications including pelvic inflammatory disease, ectopic pregnancy, sterility, and increased risk of developing cervical and ovarian cancer^4,5^. The incidence and prevalence of *C.t.* infections are rapidly rising due to a lack of long-term protective immunity and treatment failure following antibiotic therapy^2^. Understanding how *C.t.* co-opts the host cell to form its unique replicative niche is vital to developing new therapies.

All *Chlamydiae* share a biphasic developmental cycle in which the bacteria alternate between two forms: the infectious elementary body (EB) and the replicative reticulate body (RB)^3^. Upon contact with a target host cell, the EB delivers a set of pre-synthesized type III secretion system (T3SS) effector proteins into the eukaryotic cell to drive cytoskeletal rearrangements and membrane remodeling, triggering endocytosis of the pathogen^6–9^. The plasma membrane-derived compartment in which the EB resides is rapidly modified by the pathogen to form a unique replicative niche, termed the inclusion. The inclusion quickly dissociates from the endolysosomal pathway^10^, trafficking along microtubules to the peri-Golgi region^11^ where the EB differentiates into an RB and initiates replication. Following multiple rounds of division, RBs convert to EBs, and the bacteria are released by extrusion of EBs or host cell lysis to begin the infection cycle anew^12^. While it is well established that formation of an intact replicative niche is vital for *C.t.* proliferation and chlamydial disease^13,14^, how *C.t.* accomplishes such feats remains incompletely understood.

*C.t.* secreted effectors fall into two major classes: inclusion membrane proteins (Incs) and conventional T3SS (cT3SS) proteins. While substantial progress has been made in identifying and characterizing Incs^15^, comparatively little is known about cT3SS effectors. Using T3SS secretion assays in *Chlamydia*, 33 cT3SS effectors have been identified to date^16,17^. Here, we leveraged chlamydial genetics, in conjunction with large-scale unbiased affinity purification-mass spectrometry (AP-MS), to comprehensively map the host pathways targeted by cT3SS effector proteins during infection. We identified high-confidence interacting partners for 21 cT3SS proteins. Intriguingly, we show that CT584, which we have renamed *<u>C</u>hlamydia* <u>e</u>ffector <u>b</u>locking <u>n</u>uclear transport (CebN), binds to a subset of Phe-Gly (FG) nucleoporins (NUPs) and the mRNA export factor Rae1, which are host targets previously only associated with viral infection^18–22^. Our data indicate that CebN predominately localizes to the nuclear envelope in both infected and bystander cells and is necessary and sufficient to inhibit STAT1 import into the nucleus following interferon (IFN)-γ stimulation. Additionally, a CebN deficient strain of chlamydia exhibits a marked decrease in growth, further emphasizing the importance of this effector in infection. This work significantly contributes to our understanding of *C.t.* cT3SS effectors and their host targets, providing a key stepping-stone for elucidating how these effector-host interactions contribute to the pathogenesis of *C.t.* infections.

## MATERIALS AND METHODS

### Bacterial and cell culture

All proteins analyzed in this study were derived from *C.t.* L2 but as is convention in the field, we use the *C.t.* D nomenclature. *Chlamydia trachomatis* serovar L2 (LGV 434/Bu) was propagated in HeLa 229 cells (America Type Culture Collection) and EBs were purified using a gastrografin density gradient as previously described^23^. HeLa cells were propagated in RPMI 1640 medium (Thermo Fisher Scientific) supplemented with 10% Fetal Bovine Serum (Gibco) at 37°C with 5% CO_2_. A2EN cells (Kerafast) were grown in keratinocyte-serum free media (K-SFM) (Thermo Fisher Scientific) supplemented with 0.16 ng/mL epidermal growth factor, 25 μg/mL bovine pituitary extract, 0.4 mM CaCl_2_, and gentamicin at 37°C with 5% CO_2_ ^24^.

### Plasmid construction

For AP-MS, each cT3SS effector was cloned into the NotI/KpnI site of pBomb4-tet-mCherry with a FLAG-tag added to the C-terminus of each *orf* by PCR. The CebN CRISPRi gblock was cloned into pBOMBL12CRia::L2 by GenScript as previously described^25^. For the sRNA knockdown approach, a 30 nucleotide-long KD sequence that targets the sequence in the CebN 5’ UTR from -41 to -12 in respect to the start codon was cloned into pBOMB5-tet-CtrR3 by Gibson assembly to generate pBomb5-tet-CebN sRNA, as previously described^26^. CebN truncations were similarly cloned into pBomb4-tet-mCherry as FLAG-tagged fusions. For ectopic expression, CebN was cloned into the KpnI/XhoI site of pcDNA3.1-GFP. The integrity of all constructs was verified by DNA sequencing at McLab. All primers used in this study are listed in Table S1.

### Transformation of *C.t*

*C. trachomatis* serovar L2 (LGV 434/Bu) EBs were transformed as previously described^27^ with minor modifications. Briefly, plasmid DNA (5 µg), fresh *C.t.* EBs from infected host cell lysates (∼2 x10^6^ EBs), and 10 µl 5X transformation mix (50 mM Tris pH 7.4 and 250 mM CaCl_2_) were gently mixed and the final volume was adjusted to 50 µl with tissue-culture grade water. Mixtures were incubated at room temperature for 30 min. RPMI with 10% FBS (4 ml) was then added to each transformation mix and 2 ml was applied to 2 wells of a 6-well plate containing a confluent HeLa cell monolayer. Plates were centrifuged at 900 x g for 30 min and at 18h post-infection, the media was replaced with RPMI with 10% FBS containing 0.3 µg/ml penicillin G (PenG). Infectious progeny were harvested every 48h and used to infect fresh HeLa cell monolayers until viable inclusions were evident (∼2-3 passages). Expression of individual FLAG-tagged fusion proteins was induced using 10 ng/ml anhydrous tetracycline (aTc), added at time of infection, and expression was confirmed by western blotting. For pBomb4-tet-CebN CRISPRi and pBomb5-tet-CebN sRNA, 5 ng/ml anhydrous tetracycline (aTc) was added 3h post-infection and knockdown was confirmed by western blotting using 1:2000 anti-CebN antibody (kindly provided by Luís Jaime Mota^28^).

### Western blotting

For AP-MS expression verification and subsequent blots, samples were resolved either using 3-8% Tris-Acetate protein gels with Tris-Acetate SDS running buffer (proteins with MW >100kDa) or 4-12% Bis-Tris protein gels with MES running buffer (proteins with MW <100kDa). Proteins were transferred to a PVDF membrane, and following blocking with 5% milk in Tris-buffered saline with 0.1% Tween 20, were probed using anti-FLAG (Thermo Fisher Scientific, 701629), anti-GFP (Novus, NB600-597), anti-NUP54 (Proteintech, 16232-1-AP), anti-NUP153 (Novus, NBP1-81725), or anti-NUP214 (abcam, AB70497) antibodies (Table S2).

### Immunofluorescence (IF) microscopy

For visualization of CebN by stimulated emission depletion (STED) microscopy, HeLa cells were transfected with pcDNA3.1-GFP empty vector or plasmid encoding for GFP-tagged CebN. Cells were fixed with 2% formaldehyde and permeabilized with 0.1% Triton-X 100 at 24h post-transfection and stained with DAPI and NUP specific antibodies: anti-NUP54, anti-NUP153, or anti-NUP214. Images were captured on a Leica SP8 inverted microscope. Images were deconvoluted using Imaris Professional Software.

For visualization of CebN during infection, HeLa cells were infected at an MOI of 2 with WT *C.t.* or *C.t.* strains expressing a FLAG-tagged empty vector, CebN-FLAG, or TmeA-FLAG. Expression was induced for 24h using 10 ng/ml aTc added at the time of infection. Cells were fixed with 4% formaldehyde 24h post-infection and stained with DAPI (ThermoFisher Scientific), anti-FLAG (Cell Signaling, 14994T), and anti-*C.t.* HSP60 (Sigma, MABF2108). Images were captured on a Nikon A1 Confocal.

### Affinity Purification (AP)

HeLa cells, in three T175 flasks, were infected at an MOI of 2 with *C.t.* strains expressing a FLAG-tagged effector protein. Expression was induced for 24h using 10 ng/ml aTc, added at the time of infection. Four h prior to lysis, 10 µM MG132 (Millipore Sigma) was added to the media. Cells were subsequently lysed in eukaryotic lysis solution (ELS) (50 mM Tris HCl, pH 7.4, 150 mM NaCl, 1 mM EDTA, and 1% Triton-X 100) with Halt protease inhibitor cocktail (Thermo Fisher Scientific). After incubation on ice for 20 min, lysates were centrifuged at 12,000 x g for 20 min, and the supernatants were incubated with 60 µl preclearing beads (mouse IgG agarose, Millipore Sigma) for 2h at 4°C. The precleared lysate was then incubated with 30 µl FLAG beads (anti-FLAG M2 Affinity Gel, Millipore Sigma) overnight at 4°C. The beads were washed six times with ELS without detergent. For mass spectrometry, samples were stored in 50 mM ammonium bicarbonate prior to digestion and analysis as previously described^29^. For western blotting, proteins were eluted from the beads in 4X NuPAGE LDS Sample Buffer (Thermo Fisher Scientific) and boiled for 5 min.

### Mass Spectrometry (MS)

MS was performed as previously described^29^, with the following adjustments. Beads containing samples were washed with 25 mM ammonium bicarbonate and digested with 0.5 micrograms trypsin (Pierce, Thermo Fisher Scientific, MS Grade) using a CEM microwave reactor for 30 min at 55°C. Digested peptides were extracted twice using 50% acetonitrile plus 5% formic acid, lyophilized to dryness, and resuspended in 5% acetonitrile plus 0.1% formic acid. For LC/MS, samples were injected into an UltiMate 3000 UHPLC system coupled online to a Thermo Scientific Orbitrap Fusion Tribrid mass spectrometer. Peptides were separated by reversed-phase chromatography using a 50-cm MicroPac Nano C18 column (Thermo Fisher Scientific) with mobile phases of 0.1% formic acid in water and 0.1% formic acid in acetonitrile; a linear gradient from 4% to 35% Acetonitrile over the course of 45 min was employed for peptide separations. The mass spectrometer was operated in a data-dependent acquisition (DDA) mode, employing precursor scans from 300 to 1,500 m/z (120,000 resolution) followed by collision induced dissociation (CID) of the most intense precursors over a maximum cycle time of 3 s (35% NCE, 1.6 m/z isolation window, 60-s dynamic exclusion window). Raw LC-MS/MS data were converted to peak lists using Mascot Distiller 2.8 and searched against a database containing UniProt_Human and Chlamydia_trachomatis_L2434Bu using Mascot 2.8 (Matrix Science). Tryptic digestion was specified with a maximum of two missed cleavages, while peptide and fragment mass tolerances were set to 10 ppm and 0.6 Daltons, respectively. Label-free Quantitation was performed utilizing the Mascot Average method on Mascot Distiller 2.8.2.

### AP-MS analysis

Mass Spectrometry interaction STatistics (MiST) was used, as previously described ^30^, to assign a confidence score to every host protein identified from MS. Localization, Reactome pathways, biological processes, and molecular functions were determined for each host prey with a MiST^30^ score ≥0.7 using Uniprot and GeneCards. Dot plots for visualization were generated using R package ggplot2. For the protein-protein interaction network, R programming language version 4.4.1 and CORUM ^31^ were used to analyze the host proteins identified in the AP-MS and compare the dataset with previous published studies^32–34^. Cytoscape (3.10.3)^35^ was used to visualize the protein-protein interaction network.

### STAT1 and mCherry translocation assays

HeLa cells were transfected with pcDNA3.1-GFP plasmids containing empty vector, CebN, or TmeA using Lipofectamine LTX (Thermo Fisher Scientific). For mCherry-NLS translocation, 4h post-GFP-transfection the cells were transfected with mCherry-NLS (Addgene 58476). At 24h post transfection, the media was changed, with half the samples receiving normal RPMI media and half with RPMI with 100 U/ml IFN-γ for 1h. To monitor STAT1 nuclear translocation during infection, cells were infected at an MOI of 1 with or without 5 ng/ml aTc added to the media at the time of infection. At 24h post-infection, RPMI with 100 U/ml IFN-γ was added to half the samples for 1h. Transfected or infected cells were fixed with 4% formaldehyde, permeabilized with 0.1% Triton-X, blocked in 3% BSA, and stained with DAPI, anti-STAT1 (Cell Signaling 14994T), and anti-*C.t.* HSP60 antibodies (ThermoFisher Scientific MA3-023). Images were collected on Nikon Eclipse 2 microscope using the same exposure conditions between groups. For all nuclear translocation experiments (STAT1 and mCherry) nuclear signal was quantified by measuring the fluorescence intensity of STAT1/mCherry in the nucleus using Fiji with 150 transfected or infected cells per biological replicate with three replicates.

### Growth curve and inclusion measurements

HeLa or A2EN cells were infected at an MOI of 1 on ice. After 30 min, the media was changed with half the samples receiving 5 ng/ml aTc. For growth curves, samples were collected at 0 and 48 h, lysed in water and applied to fresh HeLa cell monolayers to determine the number of infectious forming units as previously described^14,36,37^. For inclusion area measurements, infected cells were fixed with 4% formaldehyde at 24h post-infection and stained using anti-*C.t.* HSP60 antibodies. Circles were drawn around the inclusions in Fiji to measure the inclusion area using 30-45 images across three coverslips per biological replicate with three replicates total.

### Statistics

Statistical analyses were performed using GraphPad Prism 10.5.0 software. Depending on the dataset, either Welch’s *t*-test or one-way ANOVAs followed by Tukey’s multiple comparisons were applied. Statistical significance was defined as P < 0.05 (*), P < 0.01 (**), and P < 0.001 (***), P < 0.0001 (****).

## RESULTS

### Identification of putative host proteins and pathways targeted by cT3SS effector proteins during *C.t.* infection

A few cT3SS effector proteins have been functionally characterized and shown to modulate diverse host cell signaling pathways^6,8,9,29,38–41^. However, the function of most of these putative virulence factors remains unknown. Affinity purification-mass spectrometry (AP-MS) has emerged as a powerful technique to comprehensively map protein-protein interactions (PPIs) between bacterial effectors or viral proteins and host proteins, yielding key mechanistic insights into how these pathogens establish their unique replicative niches^15,30,42–44^. While informative, most of these studies have been undertaken by overexpressing a single effector protein in a mammalian cell at non-physiological levels and in the absence of additional bacterial or viral factors that might promote or hinder PPIs. With the increasing ease of genetic tractability of *C.t.*, we are poised to evaluate effector-host PPIs in the context of infection.

Here, we leveraged AP-MS to systematically interrogate host interaction networks of 33 *C.t.* cT3SS effectors in the context of infection^16,17,45^. Of these, 24 were successfully expressed in *C.t.* and the remaining effector proteins were excluded from further analysis due to the inability to obtain chlamydial transformants or due to the lack of detectable expression by western blotting. Following AP-MS, only putative host target or “prey” proteins with at least 2 unique peptides that were present in at least 2 of the replicates were further pursued. CT311 and CT161 were excluded from further analysis due to lack of detection of the bait protein following AP-MS. To identify high-confidence PPIs, we analyzed the complete data set using Mass Spectrometry interaction STatistics (MiST) (Table S3), which evaluates prey reproducibility, abundance, and specificity to generate scores between 0 and 1^30^. Using a cut-off score of ≥0.7, we identified 241 putative host interacting partners for 21 cT3SS effectors. While CT144 was detected in the AP-MS, and multiple preys were identified, none of these were predicted high-confidence interactions when scored by MiST.

To further define the potential function of the *C.t.* cT3SS effector proteins, Gene Ontology (Genecards) and pathway analysis (Reactome and Uniprot) was performed for each of the 241 MiST high-confidence interactors (Table S3) (Fig. 1, S1). As shown in Supp. Fig. 1A, most of the host proteins targeted by *C.t.* effectors reside in the cytoplasm, nucleus, endoplasmic reticulum, and mitochondria; however, a few host proteins associated with the Golgi apparatus, ribosomes, cytoskeleton, and plasma membrane were also noted. To further analyze the potential function of these effector proteins, in conjunction with the above analysis, we employed Cytoscape to map PPI networks to identify effector proteins associated with multiprotein complexes or biological pathways (Fig. 1 and S1). Within this subset, we found individual cT3SS effector proteins that associate with several members of multiprotein complexes, including the proteosome (CebN), ribosome biogenesis (CT620 and CT691), tRNA processing (CT627), mRNA processing (CT386), redox homeostasis (CT053), and the nuclear pore complex (CebN) (Fig. 1). We also noted several instances of two or more cT3SS effector protein interacting with the same host protein. For example, three cT3SS effectors (CT695/TmeB, CT712, and CT392) interacted with coatomer subunit alpha (COPA) (Fig. 1), a protein required for Golgi budding and that is essential for retrograde transport and Golgi structural integrity, suggesting that these effectors may modulate COPI-dependent trafficking and Golgi architecture. Three proteins (CT736, CT691, and CT392) interacted with transferrin receptor protein 1 (TRFC), which mediates cellular iron uptake (Fig. 1). Together with prior evidence that transferrin is recruited to the inclusion in a CpoS-dependent manner^32,37,46,47^, these findings suggests that these effectors might aid in TFRC-dependent trafficking to facilitate iron acquisition. Two cT3SS effector proteins (CebN and CT053) interacted with nucleolin (NCL), a major component of the nucleolus. Most striking was the observation that CebN associates with numerous components of the nuclear pore complex (NPC). In total, we identified 68 high-confidence interactors (Table S3), many of which mediate ribonucleoprotein import, mRNA export, or transcription (Fig. 1)

**Figure 1.**
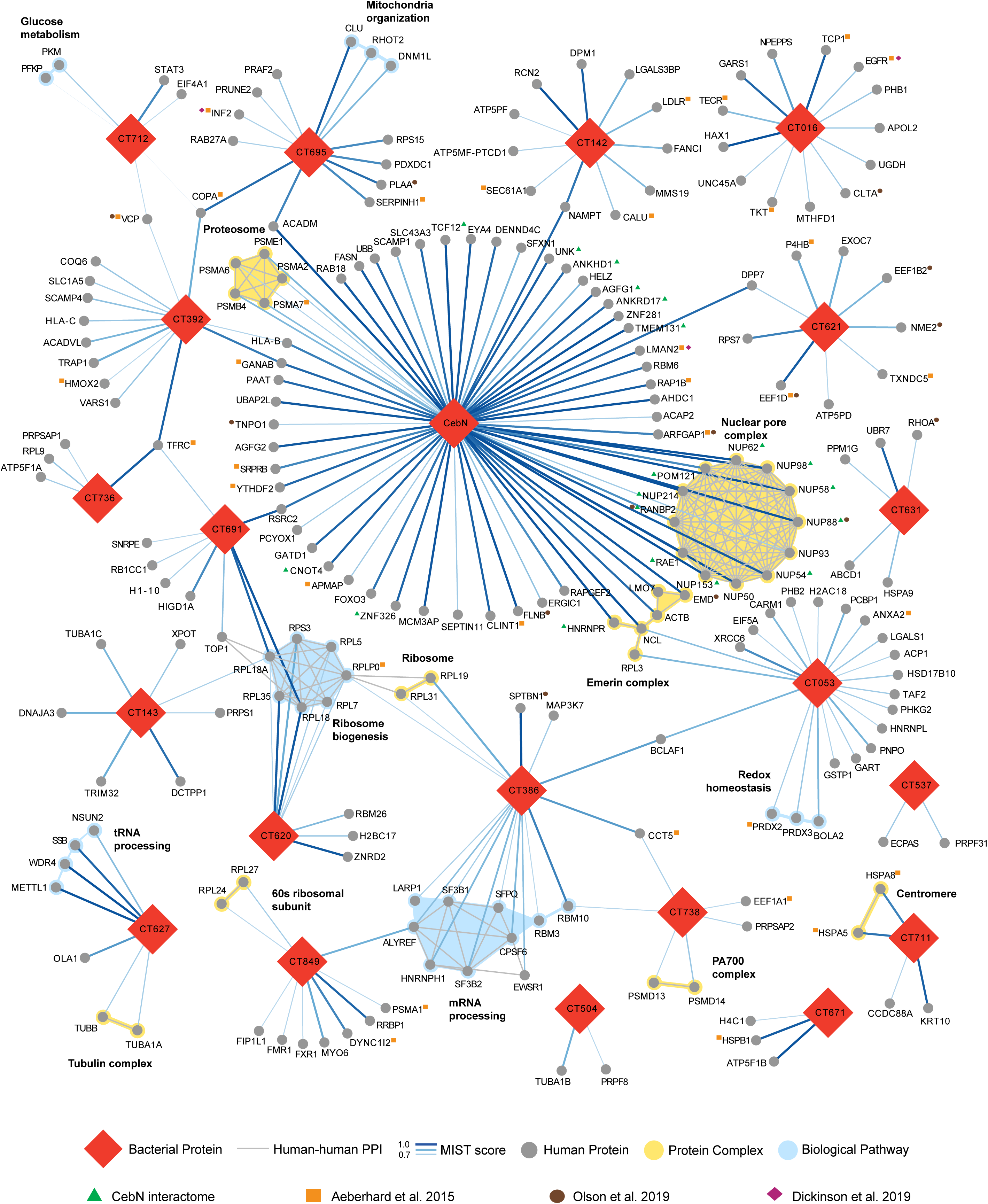
Host pathways targeted by *C.t.* secreted effector proteins identified using AP-MS. MiST was used to identify high-confidence interacting partners for each effector screened. Those with a MiST score ≥0.7 were considered significant and used for further analysis using the CORUM database to identify protein complexes targeted by each effector. An R script was used to compare the interactome to data from prior studies indicated by orange squares ^32^, brown circles ^33^, or purple diamonds^34^ next to the human protein. Red diamonds represent *C.t.* effectors; grey circles represent human proteins; yellow shading highlights protein complexes; blue shading highlights biological pathways. Human-human PPI are shown as grey lines, and bacterial-human PPI interactions as blue lines, with line thickness correlating with MiST score.

Consistent with proximity-labeling studies that identified inclusion-associated proteins^33,34^ and proteomic analyses of isolated inclusions^32^, several host factors including inverted formin 2 (IF2), COPA, transitional endoplasmic reticulum ATPase (VCP/p97), protein transport protein Sec61 subunit alpha isoform 1 (SEC61A1), epidermal growth factor receptor (EGFR), cytoplasmic dynein 1 intermediate chain 2 (DYNC1I2), amongst others, interacted with cT3SS effector proteins (Fig. 1). These concordant observations provide orthogonal validation of our AP-MS dataset and support the biological relevance of the effector-host interactome.

### Ectopically expressed CebN binds to multiple nucleoporins and Rae1

Transfected CebN-GFP predominately localized to the nuclear envelope (Fig. S2), a pattern that aligns with it binding to host proteins involved in nucleocytoplasmic transport. To confirm these interactions, and to rule out the requirement of additional bacterial proteins that contribute to the CebN infection interactome, we performed AP-MS on Strep-tagged CebN as previously described^15^. This approach identified 30 high-confidence interactors (MIST ≥ 0.7) for CebN (Table S3), of which 19 overlapped with the infection IP (Table 2, Fig. 2A). Notably, 9 nucleoporins (NUP58, NUP214, NUP98, NUP54, NUP62, NUP88, NUP153, POM121/NUP121 and RANBP2/NUP358) and the mRNA export factor Rae1 were present in both the CebN infection and transfection interactomes.

**Table 1.** Host proteins with significant MiST scores (≥0.7) identified in both infection- and transfection-based immunoprecipitations of CebN. Gene name and information regarding subcellular localization, molecular function, and biological process was obtained from Uniprot.

| Gene | Subcellular Localization | Molecular Function | Biological Process | Transfection IP MiST | Infection IP MiST |
| --- | --- | --- | --- | --- | --- |
| HNRNPR | nucleoplasm | RNA binding | mRNA processing | 1.000 | 0.992 |
| NUP58 | nuclear pore | structural component of nuclear pore complex | transport: nucleocytoplasmic | 0.995 | 0.993 |
| NUP214 | nuclear pore | structural component of nuclear pore complex | transport: nucleocytoplasmic | 0.994 | 0.994 |
| NUP98 | nuclear pore | structural component of nuclear pore complex | transport: nucleocytoplasmic | 0.993 | 0.994 |
| RAE1 | nuclear pore | RNA binding | transport: nucleocytoplasmic | 0.991 | 0.992 |
| TCF12 | nucleosome | DNA binding | transcription | 0.990 | 0.993 |
| CNOT4 | cytoplasm | transferase activity | ubiquitin-dependent protein catabolic process | 0.989 | 0.986 |
| NUP54 | nuclear pore | structural component of nuclear pore complex | transport: nucleocytoplasmic | 0.986 | 0.989 |
| ZNF326 | nuclear matrix | RNA binding | mRNA processing | 0.982 | 0.986 |
| NUP62 | nuclear pore | structural component of nuclear pore complex | transport: nucleocytoplasmic | 0.973 | 0.993 |
| NUP88 | nuclear pore | structural component of nuclear pore complex | transport: nucleocytoplasmic | 0.972 | 0.992 |
| POM121 | nuclear pore | structural component of nuclear pore complex | transport: nucleocytoplasmic | 0.971 | 0.968 |
| AGFG1 | nuclear pore | GTPase activator activity | mRNA export | 0.963 | 0.988 |
| NUP153 | nuclear pore | structural component of nuclear pore complex | transport: nucleocytoplasmic | 0.961 | 0.994 |
| UNK | cytoplasm | RNA binding | translation | 0.946 | 0.988 |
| ANKRD17 | nucleoplasm | protein binding | immune system process | 0.911 | 0.983 |
| TMEM131 | membrane | unknown | unknown | 0.866 | 0.994 |
| ANKHD1 | cytoplasm | protein binding | immune system process | 0.826 | 0.983 |
| RANBP2 | nuclear pore | ligase activity | transport: nucleocytoplasmic | 0.690 | 0.966 |

**Figure 2.**
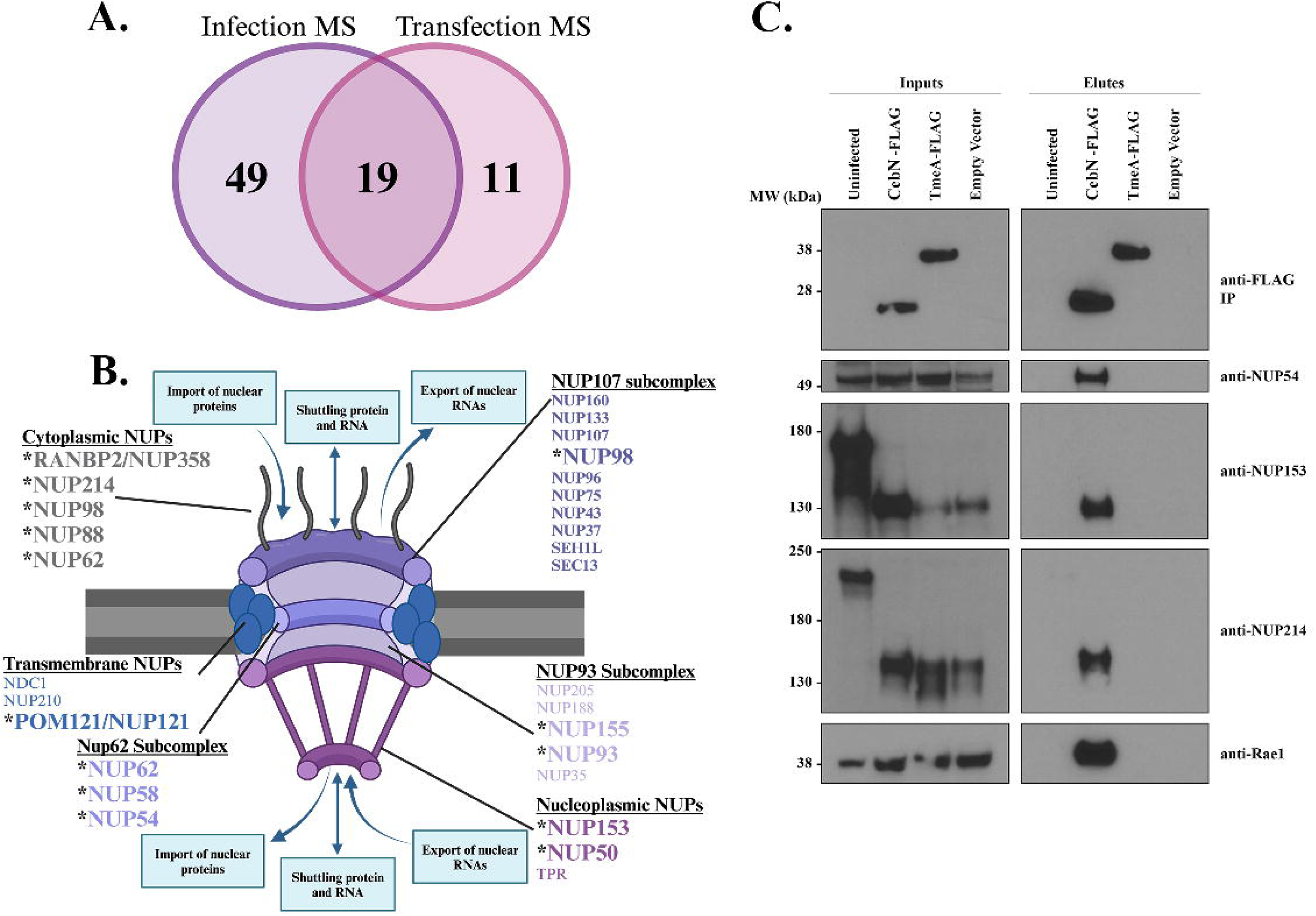
CebN interacts with multiple nucleoporins and Rae1. (A) Venn diagram comparing unique and shared hits from infection (purple) and transfection (pink) AP-MS, showing a strong overlap of identified host targets. (B) Schematic of the nuclear pore complex. Bolded and asterisked NUPs were identified as putative CebN targets using AP-MS. (C) IP of *C.t.* expressing FLAG-tagged vector, CebN, or TmeA from HeLa cells. Blots were probed with antibodies specific to selected NUPs and Rae1. Data are representative of three replicates.

Nucleoporins (NUPs) are a family of ∼30 proteins that form the nuclear pore complex (NPC) and play an important role in regulating import and export of small molecules into and out of the nucleus^48^. The NPC is organized into an inner pore ring, the nuclear and cytoplasmic rings, the nuclear basket, and the cytoplasmic filaments, each of which are enriched for select NUPs (Fig. 2B)^48^. Intriguingly, while most of the NUPs that CebN binds make up the cytoplasmic filaments, interactions with NUPs in other subcomplexes of the NPC were noted (Fig. 2B), suggesting that secreted CebN may play a broad role in modulating NPC function. Rae1 is an mRNA export factor that binds to NUP98 to aid in the transport of messenger ribonucleoprotein (mRNP) complexes through the nuclear pore complex^49^. Several viral proteins target NUPs and Rae1 to promote replication of their genomic information and to dampen the host response to infection by blocking import of important transcription factors^19–22^. To the best of our knowledge, no bacterial protein has been identified that targets NUP proteins or Rae1, making CebN an intriguing effector protein to study.

### CebN binds to and co-localizes with NUPs and Rae1

To confirm CebN binding to NUP proteins and Rae1, we immunoprecipitated FLAG tagged CebN from *C.t.* infected cells and probed with antibodies specific to NUPs and Rae1. We focused on NUP54, NUP153, and NUP214 due to their high peptide counts in the AP-MS (Table S3). NUP54, NUP153, NUP214, and Rae1 IP with CebN but not with vector or TmeA (Fig. 2C), an effector previously shown to bind to N-WASP^6,8^. Processing of NUP153 and NUP214 was noted on these blots in infected samples. We determined that CPAF, a broad-spectrum protease produced by *C.t.*, was responsible for this cleavage, as this processing was absent in lysates derived from HeLa cells infected with a CPAF mutant^50^ (Fig. S3).

To additionally confirm the interaction between CebN and NUPs, HeLa cells were transfected with GFP-CebN or GFP, fixed, and stained using anti-NUP54, NUP153, or NUP214 antibodies. Imaging by stimulated emission depletion (STED) microscopy confirmed that CebN colocalized (white) with these specific NUP proteins, whereas no colocalization was noted with GFP (Fig. 3A, B). Pearson’s correlation coefficient was calculated as a measure of colocalization, and a significant difference was found between GFP and GFP-CebN transfected cells for each individual NUP (Fig. 3B). Taken together our results indicate that the cT3SS effector protein CebN localizes to the nuclear envelope where it binds to multiple NUPs and to Rae1.

**Figure 3.**
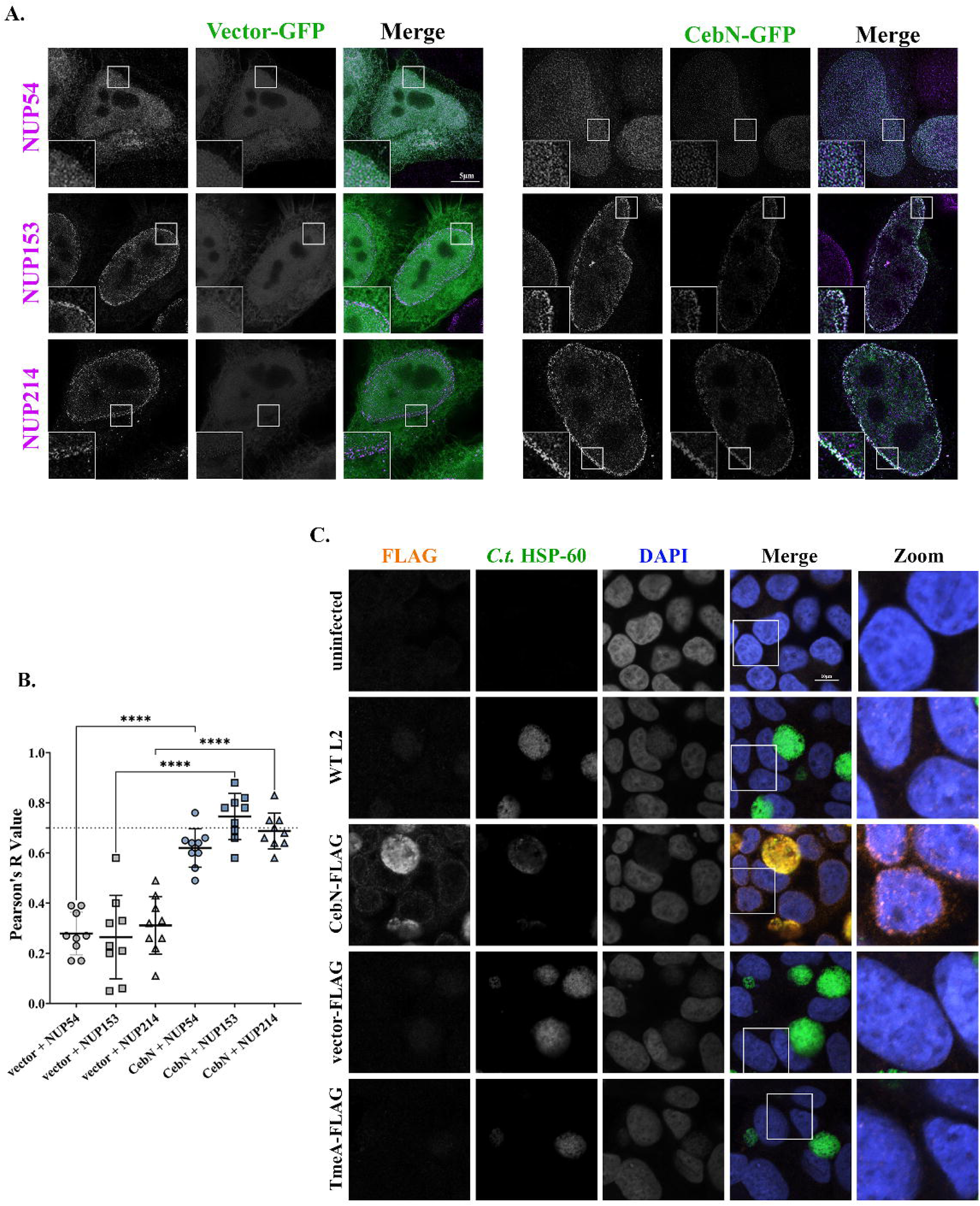
CebN localizes to the nuclear envelope during both transfection and infection conditions. (A) STED images of HeLa cells transfected with GFP-tagged empty vector or GFP- CebN (green) and stained with NUP-specific antibodies (magenta). (B) Quantification of colocalization (white) was performed in Fiji using Pearson’s correlation coefficient. Statistical significance between GFP and GFP-CebN transfected samples is shown. The graph displays individual values, the mean (black line), and standard deviation. ****P<0.0001; significance was determined using one-way ANOVA followed by Tukey’s multiple comparisons test. (C) HeLa cells uninfected or infected at an MOI of 2 with WT L2 or *C.t.* expressing FLAG-tagged vector, CebN, or TmeA (negative control). Cells were fixed with 4% formaldehyde and stained with FLAG (orange), *C.t.* HSP-60 (green) and DAPI (blue). Zoomed panels show overexposed images of the boxed region, highlighting nuclear envelopes of infected and/or bystander cells. (A, C) Data are representative of three replicates.

### CebN localizes to the nuclear envelope of infected and bystander cells

Most cT3SS effector proteins are not readily visualized by microscopy, and thus their subcellular localization is generally assessed by transfection of tagged proteins. Due to CebN’s unique localization to the nuclear envelope, we assessed CebN localization directly in *C.t.* infected cells. In line with our ectopic expression data, CebN-FLAG was found to localize to the nuclear envelope of infected cells (Fig. 3C). Intriguingly, we also observed CebN on the nuclear envelopes of bystander cells. (Fig. 3C). While one possibility is that CebN is directly translocated into neighboring cells, potentially through exosomes or tunneling nanotubes, an alternative explanation is that these signals represent effectors retained in daughter cells following mitotic division.

### The C-terminus of CebN is required for interactions with NUPs and Rae1 as well as for its localization to nuclear envelope

To delineate the region of CebN that is necessary for interaction with NUPs and Rae1, we generated 20-40 amino acid sequential truncations from the C-terminus of the 183 amino acid protein and expressed these truncations as FLAG-tagged constructs in *C.t.* Immunoprecipitation of these truncations, followed by subsequent western blotting, showed that the C-terminal 23 amino acids of CebN are necessary for interactions with NUP54, NUP153, NUP214, and Rae1 (Fig. 4A). We further confirmed the importance of this region by performing confocal microscopy on HeLa cells infected with *C.t.* strains expressing the CebN FLAG-tagged deletion constructs (Fig 4B). As noted above, full length CebN-FLAG localizes to the nuclear envelope of infected cells, as well as the nuclear envelopes of bystander cells (Fig. 4B). However, none of the truncated versions of CebN localized to the nuclear envelope in infected or bystander cells (Fig. 4B). In total, the C-terminus of CebN is necessary for its interaction with NUPs and Rae1 and for its localization to the nuclear envelope in infected and bystander cells.

**Figure 4.**
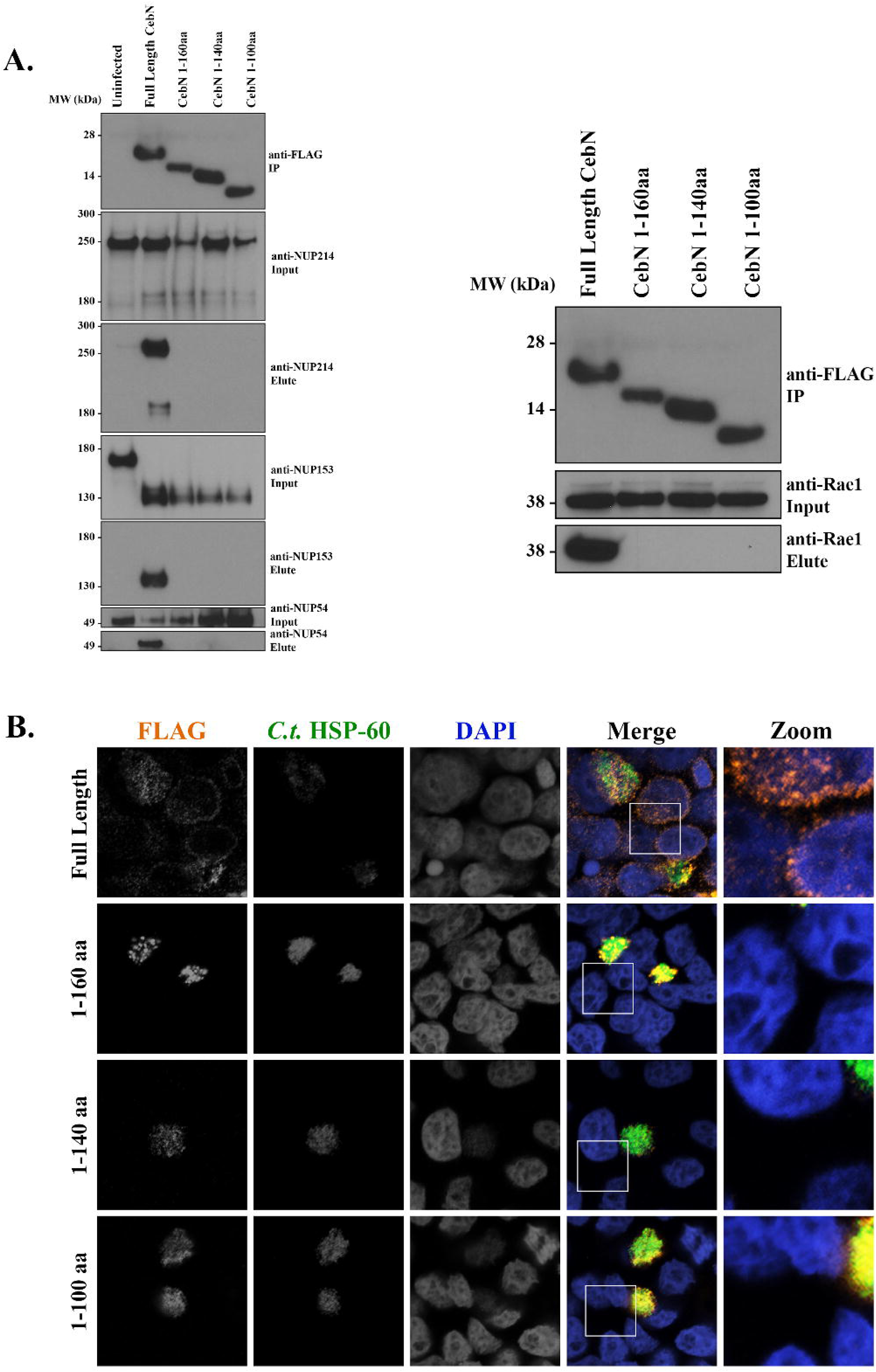
The C-terminus of CebN mediates interactions with NUPs and Rae1 and is required for nuclear envelope localization. (A) Lysates from HeLa cells infected for 24 h with *C.t.* expressing the indicated C-terminally FLAG-tagged CebN constructs were immunoprecipitated on FLAG beads and immunoblotted with antibodies specific to selected NUPs and Rae1. (B) HeLa cells infected for 24 h with *C.t.* expressing the indicated C-terminally FLAG-tagged CebN constructs were fixed with 4% formaldehyde and stained with antibodies to FLAG (orange), *C.t.* HSP-60 (green, to visualize bacteria), and with DAPI (blue). Zoomed panels show overexposures of the boxed region highlighting nuclear envelopes of infected and/or bystander cells. (A-B) Data are representative of at least two replicates.

### CebN is important for chlamydial replication *in vtiro*

To determine the importance of CebN during *Chlamydia* infection, we generated two conditional CebN mutants, using CRISPRi and sRNA systems^25,26,51^, in which expression of CebN is repressed upon induction with anhydrous tetracycline. Knockdown of CebN for both mutant strains was confirmed by western blotting (Fig. 5A). CRISPRi was implemented using a deactivated Cas 12 enzyme that binds 186bp downstream of the *cebN* start site, thereby blocking effective transcription of the gene. In parallel, the sRNA system employed engineered small RNAs that base-pair with *cebN* mRNA transcripts to inhibit translation of CebN protein. Together, these complementary approaches enable conditional knockdown of CebN at both the transcriptional and translational levels. In both HeLa and A2EN cells, both mutants exhibited significantly reduced growth upon aTc induction compared to uninduced or wild-type infections (Fig. 5B). The growth defect was further reflected by a decrease in inclusion size in both cell types (Fig. 5C). Collectively, these results demonstrate that CebN is required for efficient bacterial replication and inclusion development.

**Figure 5.**
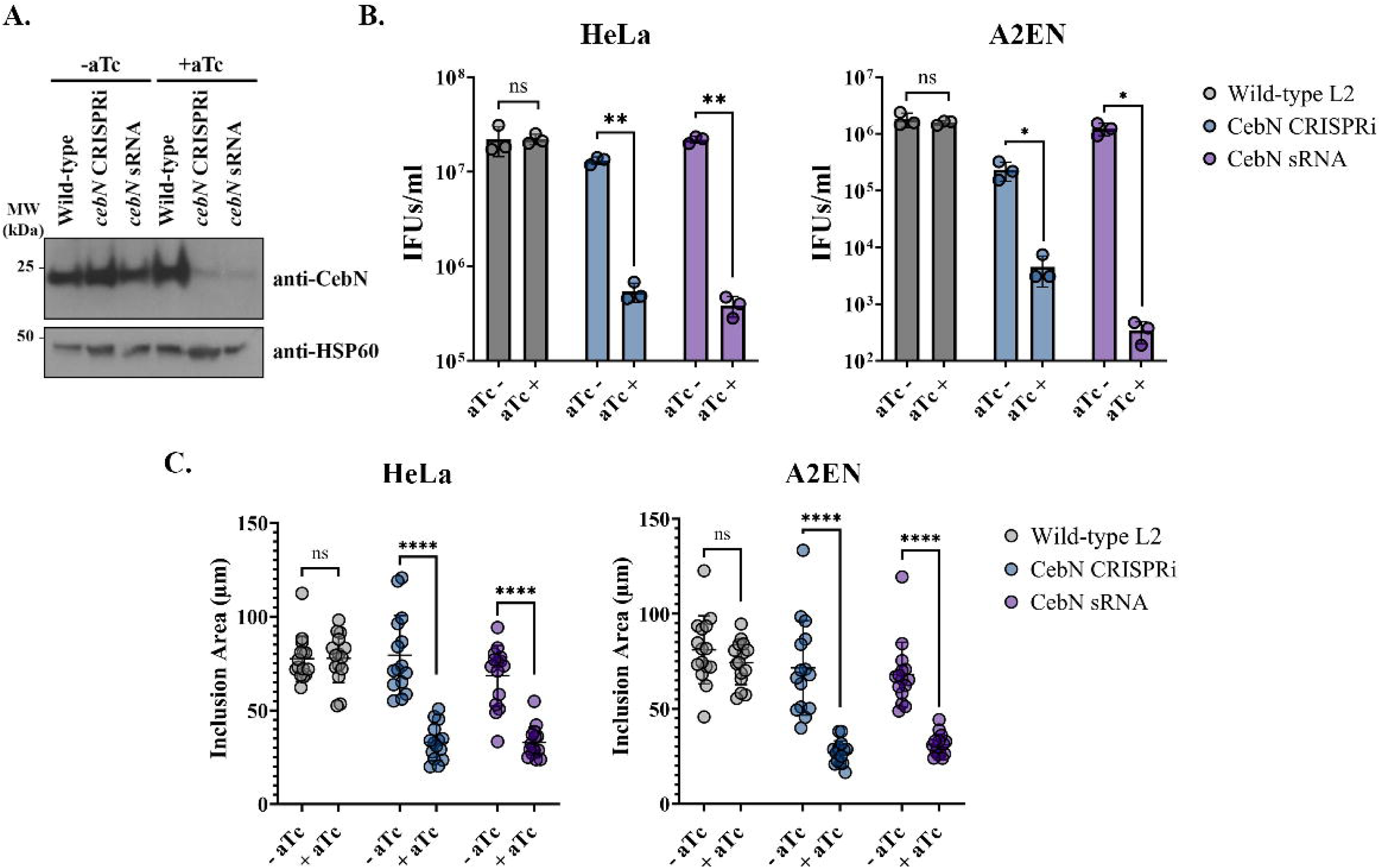
CebN is important for intracellular replication and inclusion development. (A) Western blot confirmation of CebN knockdown in lysates from HeLa cells infected with CRISPRi and sRNA chlamydial strains with or without aTc induction. (B) Quantification of infectious progeny at 48 h postinfection in HeLa (left) or A2EN (right) cells infected with WT (grey), CebN CRISPRi mutant (blue), or CebN sRNA mutant (purple). (C) Quantification of inclusion areas (µm^2^) in HeLa (left) and A2EN (right) cells infected with WT (grey), CebN CRISPRi mutant (blue), or CebN sRNA mutant (purple) *C.t.* strains. (B-C) Statistical significance was determined using Welch’s t-test. *P<0.05, **P<0.01, ****P<0.0001. Error bars are standard deviation. (A- C) Data are representative of three replicates.

### CebN attenuates STAT1 import into the nucleus following interferon-γ stimulation

Viral proteins from HIV, SARS-CoV-2, Kaposi’s sarcoma-associated herpesvirus, and vesicular stomatitis virus interact with and remodel the nuclear pore complex to modulate nuclear import of transcription factors required for the antiviral response^19,20,22,52–58^. Similarly, *C.t.* attenuates STAT1 nuclear import following IFN-γ stimulation^59^. We hypothesized that CebN interactions with NUPs and Rae1 could underlie this perturbation. To test this, HeLa cells were transfected with GFP-CebN, GFP-empty vector, or GFP-TmeA, treated with IFN-γ, and imaged by confocal immunofluorescence microscopy using an anti-STAT1 antibody (Fig. 6A). We observed a significant reduction in the proportion of cells with nuclear STAT1 in CebN-transfected cells compared to those transfected with empty vector or TmeA (Fig. 6B), indicating that CebN is sufficient to impair STAT1 nuclear translocation.

**Figure 6.**
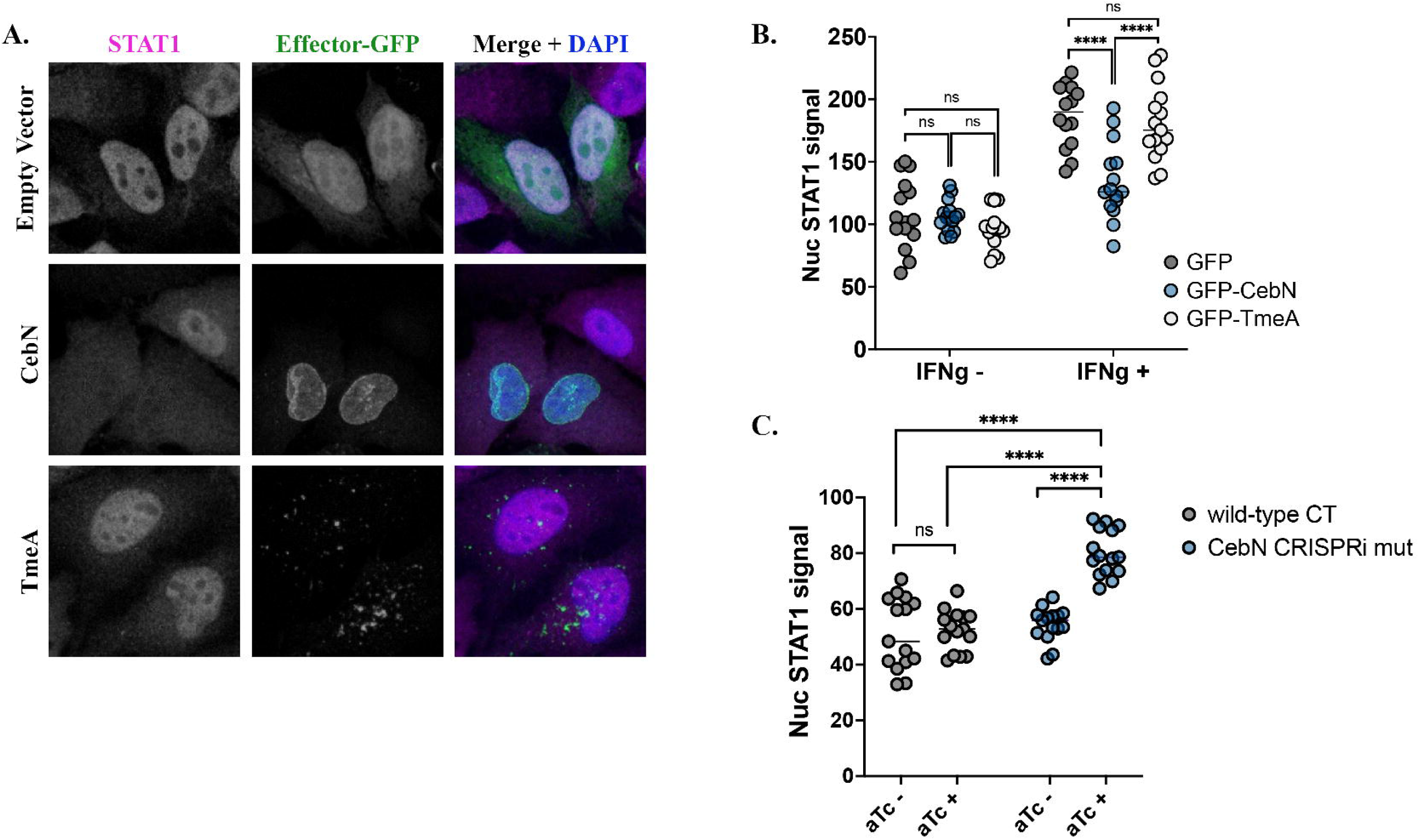
CebN is necessary and sufficient to perturb IFN-γ-stimulated STAT1 nuclear translocation. (A) HeLa cells were transfected with GFP-tagged vector, GFP-CebN, or GFP-TmeA (green) and treated with 600U/ml IFN-γ. Cells were fixed and stained with anti-STAT1 (magenta) antibody and DAPI to demark the nucleus (blue). (B, C) Translocation of STAT1 into the nucleus of transfected (B) or infected cells (C) was quantified in relative fluorescence units (RFUs) from 150 cells per biological replicate. Error bars represent standard deviation. Statistical significance was determined using one-way ANOVA followed by Tukey’s multiple comparisons test. ****P<0.0001. (A-C) Data are representative of three replicates.

To test whether CebN is necessary for this process during infection, HeLa cells were infected with WT or CebN CRISPRi mutant strain, treated with IFN-γ, stained with anti-STAT1 antibody, and nuclear STAT1 signal was quantified. We observed a significant increase in STAT1 nuclear localization when the CebN mutant strain was induced with aTc, demonstrating that CebN is required to block STAT1 nuclear translocation (Fig. 6C).

To assess whether this effect was due to a general block in nuclear import, we used a nuclear localization signal (NLS)-tagged mCherry reporter plasmid. In HeLa cells co-transfected with mCherry-NLS and GFP-empty vector, GFP-CebN, or GFP-TmeA, we detected no significant differences in nuclear mCherry signal, suggesting that CebN selectively interferes with import rather than imparting a general blockade (Fig. S4). Taken together, our results suggest that CebN, through interactions with nucleoporins and Rae1, plays a key role in dampening the host response to *C.t.* infection by perturbing nuclear translocation of transcriptional regulators of the host innate immune response.

## DISCUSSION

In this study, we combined *C.t.* genetics with AP-MS to generate the first cT3SS effector-host interactome. Our approach successfully identified high confidence interacting host partners for 21 of the 36 uncharacterized cT3SS effectors. Our work is especially valuable as it not only begins to build a compendium of PPIs during active infection but also provides a launch point for detailed mechanistic characterization of these effector proteins. Importantly, screens such as AP-MS, can reveal novel pathways targeted by intracellular bacteria^42–44^. Here we discovered that CebN targets NUPs to impair STAT1 nuclear import, revealing a potential mechanism by which *C.t.* dampens the host innate immune response to survive intracellularly. Altogether, these studies enhance our understanding of how obligate intracellular pathogens remodel the host to form their unique replicative niches.

One of the most striking findings from our AP-MS screen was the interaction between CebN and multiple nucleoporins and Rae1. While our study is, to the best of our knowledge, the first time a bacterial effector has been shown to interact with host nucleoporin proteins, several viral proteins have been identified that co-opt NUPs and Rae1. ORF6 of SARS-CoV-2, ORF10 of Kaposi’s sarcoma-associated herpesvirus, and M protein of vesicular stomatitis virus all bind to the NUP98-Rae1 complex, whereas the HIV-1 capsid binds to multiple nucleoporins leading to altered NUP expression and localization^19,20,22,52–58^. While HIV-1 manipulates NUPs to facilitate viral import and integration of its genome into the host genome^60^, other viral proteins interact with NUPs to disrupt nucleocytoplasmic transport of key transcription factors such as STAT1. During SARS-CoV-2 infection, ORF6-Rae1-NUP98 interactions block STAT1 nuclear import and mRNA export, resulting in a significantly diminished host response^18,20,22,57^. The ability of CebN to perturb STAT1 import during *C.t.* infection, along with our observation that it binds NUPs-Rae1, suggests it may subvert host defense mechanisms in a manner reminiscent of viral proteins such as ORF6. However, given that nucleoporins and Rae1 regulate a broad range of cellular processes including mRNA export, protein import, transcriptional regulation, and chromatin organization^48^, it remains possible that CebN’s primary and physiologically relevant function involves one or more of these other pathways.

Similar to how viruses dampen the immune response, *C.t.* has been shown to antagonize interferon pathways^61–66^. IFN production activates the JAK-STAT signaling pathway, leading to phosphorylation and homodimerization of STAT1, which is imported into the nucleus by karyopherin alpha 1 and karyopherin beta 1 heterodimers^48^. Once in the nucleus, the STAT1 homodimer complex binds to gamma-activated site promoter elements to drive expression of a subset of ISGs meant to impede the infection. Several studies have shown that following prolonged IFN-γ stimulation, nuclear translocation of STAT1 is reduced^59,62^ and inhibition of the JAK-STAT pathway correlates with lower mRNA and protein levels of key interferon response elements in infected cells compared to uninfected controls^67^. Furthermore, this difference was dependent on *de novo C.t.* protein synthesis, supporting the role of a *C.t.* effector protein in this process^67^. Our new data demonstrate that CebN is necessary and sufficient to decrease STAT1 import immediately following IFN-γ stimulation (1 h). Importantly, we show that this inhibition is not global, as CebN does not block import of NLS-tagged mCherry. These findings suggest that CebN impairs nuclear import of some factors but does not impart a total nuclear blockade. Consistent with this, prior work by Walsh et al.^65^ demonstrated that *C.t.* mutants lacking the Inc GarD are highly susceptible to IFN-γ-mediated killing due to the activity of the interferon-stimulated gene RNF213, and that RNF213 is induced at comparable levels in uninfected and infected cells. Thus, while CebN limits STAT1 import, this may not necessarily result in decreased expression of all ISGs, likely reflecting redundancy in IFN-γ-activated host defense pathways. Determining the extent to which CebN-mediated inhibition of STAT1 impacts specific ISGs is an area of future investigation.

Lining the NPC are intrinsically disordered NUPs that harbor numerous Phe-Gly (FG) repeats separated by a hydrophilic spacer of 5-30 amino acids^48^. Movement of large cargo across the NPC requires highly specific interactions between these so-called FG-NUPs and transporters of the karyopherin family, which enables entry and rapid diffusion of the cargo-transporter complex through the NPC. Of the 11 NUPs identified as putative binding partners of CebN, 9 are classified as FG-NUPs (Table 2, S3). Recognition of FG motifs within these select NUPs might explain how CebN binds to multiple nucleoporins and is able to selectively inhibit nuclear import.

Crystallization of CebN revealed an N-terminal four-helix bundle (α1-4), followed by a three-stranded antiparallel β-sheets (β1-3)^68^, and a C-terminal kinked antiparallel pair of α-helices^68^. AlphaFold modeling predicts that the C-terminus of CebN harbors a coiled-coil domain (amino acids 153-181). Truncation of the last 23 amino acids of CebN would disrupt this predicted coiled-coil, thus abrogating binding to NUPs and Rae1. While analysis of CebN did not identify motifs known to be required for interactions with NUPs or Rae1, new motifs are constantly being discovered, and it is possible that CebN possesses a previously undefined motif. Short linear motifs (SLiMs), short 3-15 amino acid motifs often embedded in intrinsically disordered or coiled-coil regions, serve as binding interfaces for structured partners and are abundant in the human proteome as well as in viral and, less frequently, bacterial effectors^69,70^.

Ectopically expressed CebN appeared to concentrate at the nuclear envelope, and we observed a similar localization in infected cells and unexpectedly also in neighboring bystander cells. This unique localization in apparently uninfected bystander cells has only ever been reported once before with the *C. psittaci* cT3SS effector SINC, which similarly targets the nuclear envelope through interactions with lamins^71^. How effectors reach bystander cells remains unknown, but proposed mechanisms include packaging into exosomes for release at the cell surface, tunneling nanotubes, or effectors leftover during cell division prior to inclusion segregation into one daughter cell. Prior work with *Mycobacterium tuberculosis* has revealed that mycobacterial proteins are packaged into vesicles and released via calcium-regulated lysosomal exocytosis. These proteins are then trafficked to uninfected bystander cells^72^. Tunneling nanotubes, on the other hand, are transient cellular connections that play a role in cell- to-cell communication and facilitate exchange of molecules between cells. Analogous to viruses, previous work has shown that *C.t.* may spread cell-to-cell by nanotubules^73^. Thus, it is conceivable that effector proteins may also be transmitted to adjacent cells via this mechanism. If an infected cell undergoes cell division, the inclusion is partitioned into one cell, leaving the other daughter cell “uninfected^74^.” It is intriguing to consider that secreted effector proteins may be left behind and continue to function in the absence of infection. Future work will involve delineating the exact mechanism by which *C.t.* is able to transport effector proteins into neighboring, uninfected cells and whether effectors besides CebN and SINC can access adjacent cells.

As with all screens, false positives and negatives can result. To add rigor to our AP-MS data set analysis, we employed MiST, which combines metrics of reproducibility, specificity, and abundance across the entire data set to identify putative host binding partners more accurately and stringently. Using this technique, we identified high confidence targets for 21 of the secreted effectors tested herein. In our study, we sought to define PPIs for all the previously uncharacterized cT3SS effector proteins and included CT695 (TmeB) as at the onset of this study TmeB had no function ascribed to it. Recent work has since shown that ectopically expressed TmeB targets the ARP2/3 complex^75^. While we did not find components of the ARP2/3 complex in our infection AP-MS, we did identify an actin-binding protein, inverted formin 2 (INF2). INF2 belongs to the formin family of proteins, which function to both polymerize and depolymerize actin filaments^76^. Formins and the ARP2/3 complex act in parallel to regulate the actin cytoskeleton^77^. Differences in experimental set-up between transfection of TmeB-FLAG and infection of a *C.t.* strain expressing TmeB-FLAG could contribute to these differences in identified putative host binding partners. Additionally, our timepoint of 24 hours post infection may correlate with functions of TmeB beyond invasion.

By combining chlamydial genetics with large-scale screens, we have begun to define the compendium of putative host proteins and pathways targeted by secreted *C.t.* effector proteins during infection. Our analysis uncovered a wealth of unique high-confidence host interactors, laying the groundwork for detailed mechanistic characterization of cT3SS effector proteins. Importantly, our approach has identified targets previously not associated with bacterial infection, including nucleoporins, which are commonly targeted by viral proteins to modulate host defenses^19–22^. We propose a model in which secretion of CebN leads to inhibition of STAT1 nuclear translocation through interactions with NUPs and Rae1 (Fig. 7), a mechanism that may alter host cell transcription and contribute to suppression of the innate immune response in infected and uninfected bystander cells. Moreover, the ability of CebN to translocate to bystander cells may provide a mechanism by which *C.t.* can prime nearby cells for infection. Understanding how CebN co-opts NUPs and Rae1 will not only advance our understanding of how *C.t.* establishes a persistent infection despite a robust host response but may also identify druggable targets applicable to both bacterial and viral infections.

**Figure 7.**
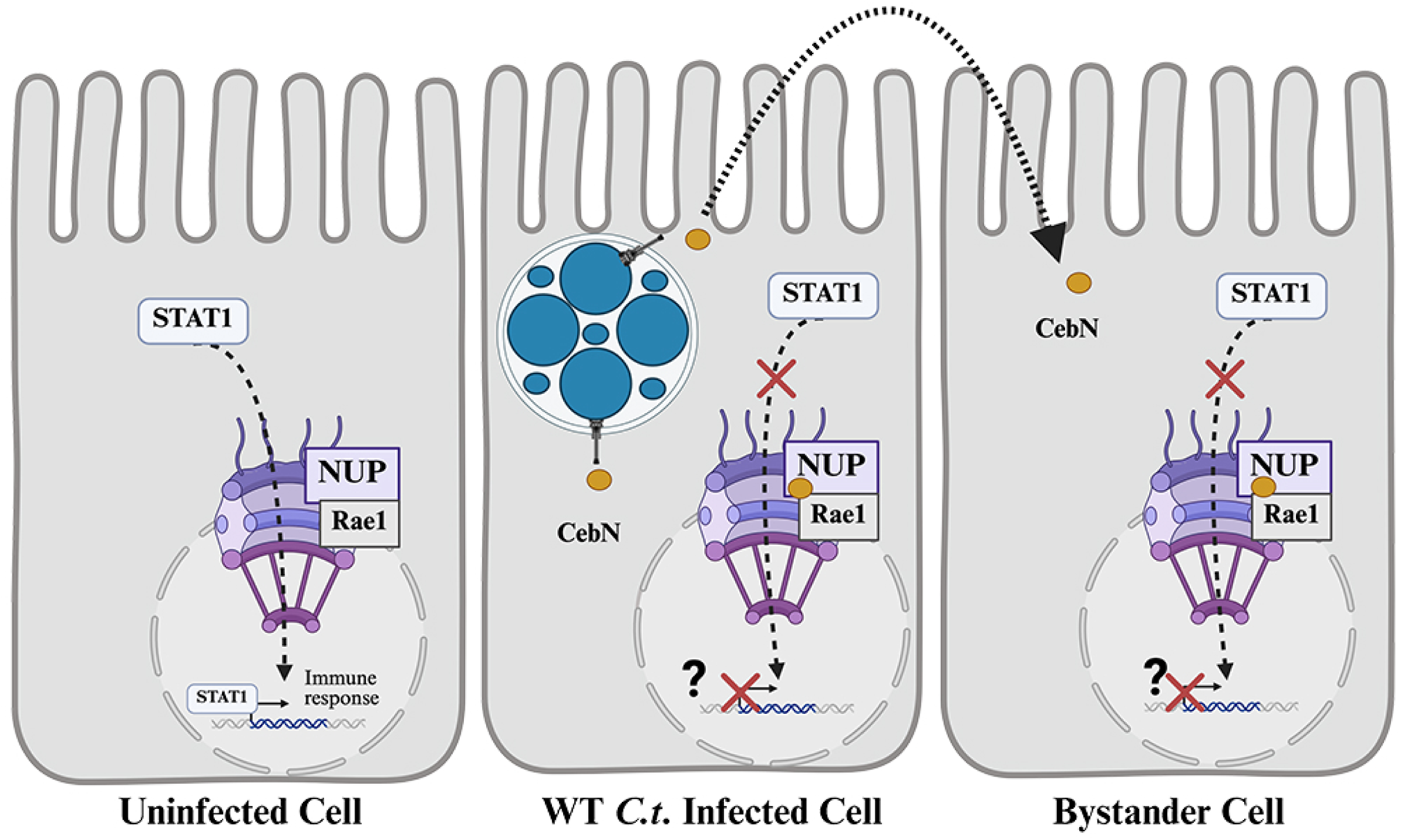
Model for CebN-mediated inhibition of STAT1 nuclear translocation. In uninfected cells, IFN-γ stimulation triggers STAT1 translocation to the nucleus through the nuclear pore complex, where STAT1 binds to gamma-activated site (GAS) promoter elements to drive expression of a subset of interferon-stimulated genes that restrict infection. In *C.t.* infected cells, STAT1 translocation is blocked by CebN, which interacts with NUPs and Rae1. We hypothesize this interaction reduces the transcriptional activity of STAT1. In addition, CebN can be translocated into neighboring cells, where it may alter nuclear import and/or export pathways to prime these cells for subsequent infection.

## Supporting information

S1 Table Primers

S2 Antibodies

S2 Mist

## Acknowledgements

We thank Dr. Raphael Valdivia for kindly sharing the *cpa*::*cat* mutant and Dr. Luís Jaime Mota for the CebN antibody. We thank Dr. Scot Ouellette for technical assistance with generation of the CRISPRi mutant. Thanks to Dr. Dominique Limoli for assistance with troubleshooting CebN staining during infection. We would like to thank Jawad Haider for excellent technical assistance and Steven Huang for critical review of this manuscript. We acknowledge grant support from the NIH (M.M.W.: R01 AI150812, R01 AI155434, R61 AI179999; J.E.: R01 AI073770, R01 AI105561; N.J.K: P50 GM082250; PO1 AI090935, PO1 090935, P50 GM081879, PO1 091575, U19 AI106754, U54AI081680, DARPA-10-93-Prophecy-PA-008), University of Iowa Training in Mechanisms of Parasitism T32 AI007511 (B.S. and X.T.), and the University of California San Francisco Pathogenesis and Host Defense T32 AI060537 (K.M.M). We also acknowledge support from the University of Iowa Stead Family Scholars Program (M.M.W.), the University of Iowa Stinski fellowship (S.E.A), UCSF Program for Breakthrough in Biomedical Research (J.E. and N.K.), and the American Heart Association (K.M.M.). The Krogan Laboratory has received research support from Vir Biotechnology, F. Hoffmann-La Roche, and Rezo Therapeutics. Nevan Krogan has a financially compensated consulting agreement with Maze Therapeutics. He is the President and is on the Board of Directors of Rezo Therapeutics, and he is a shareholder in Tenaya Therapeutics, Maze Therapeutics, Rezo Therapeutics, GEn1E Lifesciences, and Interline Therapeutics.

## REFERENCES

1. Andersen, S.E., Bulman, L.M., Steiert, B., Faris, R., and Weber, M.M. (2021). Got mutants? How advances in chlamydial genetics have furthered the study of effector proteins. Pathog Dis 79, ftaa078-. 10.1093/femspd/ftaa078.

2. 2. Maza, L.M. de la, Darville, T.L., and Pal, S. (2021). Chlamydia trachomatis vaccines for genital infections: where are we and how far is there to go? Expert Rev Vaccines 20, 421–435. 10.1080/14760584.2021.1899817.

3. Elwell, C., Mirrashidi, K., and Engel, J. (2016). Chlamydia cell biology and pathogenesis. Nat Rev Microbiol 14, 385–400. 10.1038/nrmicro.2016.30.

4. Das, M. (2018). Chlamydia infection and ovarian cancer risk. Lancet Oncol 19, e338. 10.1016/s1470-2045(18)30421-2.

5. Koskela, P., Anttila, T., Bjørge, T., Brunsvig, A., Dillner, J., Hakama, M., Hakulinen, T., Jellum, E., Lehtinen, M., Lenner, P., et al. (2000). Chlamydia trachomatis infection as a risk factor for invasive cervical cancer. Int. J. Cancer 85, 35–39. 10.1002/(sici)1097-0215(20000101)85:1<35::aid-ijc6>3.0.co;2-a.

6. Faris, R., McCullough, A., Andersen, S.E., Moninger, T.O., and Weber, M.M. (2020). The Chlamydia trachomatis secreted effector TmeA hijacks the N-WASP-ARP2/3 actin remodeling axis to facilitate cellular invasion. Plos Pathog 16, e1008878. 10.1371/journal.ppat.1008878.

7. Keb, G., Hayman, R., and Fields, K.A. (2018). Floxed-Cassette Allelic Exchange Mutagenesis Enables Markerless Gene Deletion in Chlamydia trachomatis and Can Reverse Cassette-Induced Polar Effects. J. Bacteriol. 200. 10.1128/jb.00479-18.

8. Keb, G., Ferrell, J., Scanlon, K.R., Jewett, T.J., and Fields, K.A. (2021). Chlamydia trachomatis TmeA Directly Activates N-WASP To Promote Actin Polymerization and Functions Synergistically with TarP during Invasion. Mbio 12, e02861–20. 10.1128/mbio.02861-20.

9. Ghosh, S., Ruelke, E.A., Ferrell, J.C., Bodero, M.D., Fields, K.A., and Jewett, T.J. (2020). Fluorescence-Reported Allelic Exchange Mutagenesis-Mediated Gene Deletion Indicates a Requirement for Chlamydia trachomatis Tarp during In Vivo Infectivity and Reveals a Specific Role for the C Terminus during Cellular Invasion. Infect. Immun. 88, 423–428. 10.1128/iai.00841-19.

10. Scidmore, M.A., Fischer, E.R., and Hackstadt, T. (2003). Restricted Fusion of Chlamydia trachomatis Vesicles with Endocytic Compartments during the Initial Stages of Infection. Infect. Immun. 71, 973–984. 10.1128/iai.71.2.973-984.2003.

11. Grieshaber, S.S., Grieshaber, N.A., and Hackstadt, T. (2003). Chlamydia trachomatis uses host cell dynein to traffic to the microtubule-organizing center in a p50 dynamitin-independent process. J Cell Sci 116, 3793–3802. 10.1242/jcs.00695.

12. Hybiske, K., and Stephens, R.S. (2007). Mechanisms of host cell exit by the intracellular bacterium Chlamydia. Proc National Acad Sci 104, 11430–11435. 10.1073/pnas.0703218104.

13. Sixt, B.S., Bastidas, R.J., Finethy, R., Baxter, R.M., Carpenter, V.K., Kroemer, G., Coers, J., and Valdivia, R.H. (2017). The Chlamydia trachomatis Inclusion Membrane Protein CpoS Counteracts STING-Mediated Cellular Surveillance and Suicide Programs. Cell Host Microbe 21, 113–121. 10.1016/j.chom.2016.12.002.

14. Weber, M.M., Lam, J.L., Dooley, C.A., Noriea, N.F., Hansen, B.T., Hoyt, F.H., Carmody, A.B., Sturdevant, G.L., and Hackstadt, T. (2017). Absence of Specific Chlamydia trachomatis Inclusion Membrane Proteins Triggers Premature Inclusion Membrane Lysis and Host Cell Death. Cell Reports 19, 1406–1417. 10.1016/j.celrep.2017.04.058.

15. 15. Mirrashidi, K.M., Elwell, C.A., Verschueren, E., Johnson, J.R., Frando, A., Von Dollen, J., Rosenberg, O., Gulbahce, N., Jang, G., Johnson, T., et al. (2015). Global Mapping of the Inc- Human Interactome Reveals that Retromer Restricts Chlamydia Infection. Cell Host Microbe 18, 109–121. 10.1016/j.chom.2015.06.004.

16. McCaslin, P.N., Andersen, S.E., Icardi, C.M., Faris, R., Steiert, B., Smith, P., Haider, J., and Weber, M.M. (2023). Identification and Preliminary Characterization of Novel Type III Secreted Effector Proteins in Chlamydia trachomatis. Infect. Immun., e0049122. 10.1128/iai.00491-22.

17. Faris, R., Koch, R., McCaslin, P., Challagundla, N., Steiert, B., Andersen, S.E., Smith, P., Jabeena, C.A., Yau, P., Rudel, T., et al. (2025). The Chlamydia trachomatis secreted effector protein CT181 binds to Mcl-1 to prolong neutrophil survival. bioRxiv, 2025.03.16.643443. 10.1101/2025.03.16.643443.

18. Gao, X., Tian, H., Zhu, K., Li, Q., Hao, W., Wang, L., Qin, B., Deng, H., and Cui, S. (2022). Structural basis for Sarbecovirus ORF6 mediated blockage of nucleocytoplasmic transport. Nat. Commun. 13, 4782. 10.1038/s41467-022-32489-5.

19. Kane, M., Rebensburg, S.V., Takata, M.A., Zang, T.M., Yamashita, M., Kvaratskhelia, M., and Bieniasz, P.D. (2018). Nuclear pore heterogeneity influences HIV-1 infection and the antiviral activity of MX2. eLife 7, e35738. 10.7554/elife.35738.

20. Kehrer, T., Cupic, A., Ye, C., Yildiz, S., Bouhaddou, M., Crossland, N.A., Barrall, E.A., Cohen, P., Tseng, A., Çağatay, T., et al. (2023). Impact of SARS-CoV-2 ORF6 and its variant polymorphisms on host responses and viral pathogenesis. Cell Host Microbe 31, 1668–1684.e12. 10.1016/j.chom.2023.08.003.

21. Hall, R., Guedán, A., Yap, M.W., Young, G.R., Harvey, R., Stoye, J.P., and Bishop, K.N. (2022). SARS-CoV-2 ORF6 disrupts innate immune signalling by inhibiting cellular mRNA export. PLoS Pathog. 18, e1010349. 10.1371/journal.ppat.1010349.

22. Addetia, A., Lieberman, N.A.P., Phung, Q., Hsiang, T.-Y., Xie, H., Roychoudhury, P., Shrestha, L., Loprieno, M.A., Huang, M.-L., Gale, M., et al. (2021). SARS-CoV-2 ORF6 Disrupts Bidirectional Nucleocytoplasmic Transport through Interactions with Rae1 and Nup98. mBio 12, e00065–21. 10.1128/mbio.00065-21.

23. Faris, R., and Weber, M. (2019). Propagation and Purification of Chlamydia trachomatis Serovar L2 Transformants and Mutants. Bio-protocol 9, e3459. 10.21769/bioprotoc.3459.

24. Buckner, L.R., Schust, D.J., Ding, J., Nagamatsu, T., Beatty, W., Chang, T.L., Greene, S.J., Lewis, M.E., Ruiz, B., Holman, S.L., et al. (2011). Innate immune mediator profiles and their regulation in a novel polarized immortalized epithelial cell model derived from human endocervix. J Reprod Immunol 92, 8–20. 10.1016/j.jri.2011.08.002.

25. Ouellette, S.P., Blay, E.A., Hatch, N.D., and Fisher-Marvin, L.A. (2021). CRISPR Interference To Inducibly Repress Gene Expression in Chlamydia trachomatis. Infect. Immun. 89, e00108–21. 10.1128/iai.00108-21.

26. Ehses, J., Wang, K., Densi, A., Ramirez, C., Tan, M., and Sütterlin, C. (2024). Development of an sRNA-mediated conditional knockdown system for Chlamydia trachomatis. mBio, e0254524. 10.1128/mbio.02545-24.

27. Weber, M.M., and Faris, R. (2019). Mutagenesis of Chlamydia trachomatis Using TargeTron. Methods Mol. Biol. 2042, 165–184. 10.1007/978-1-4939-9694-0_12.

28. Pais, S.V., Milho, C., Almeida, F., and Mota, L.J. (2013). Identification of Novel Type III Secretion Chaperone-Substrate Complexes of Chlamydia trachomatis. Plos One 8, e56292. 10.1371/journal.pone.0056292.

29. Steiert, B., Icardi, C.M., Faris, R., McCaslin, P.N., Smith, P., Klingelhutz, A.J., Yau, P.M., and Weber, M.M. (2023). The Chlamydia trachomatis type III-secreted effector protein CteG induces centrosome amplification through interactions with centrin-2. Proc National Acad Sci 120, e2303487120. 10.1073/pnas.2303487120.

30. Jäger, S., Cimermancic, P., Gulbahce, N., Johnson, J.R., McGovern, K.E., Clarke, S.C., Shales, M., Mercenne, G., Pache, L., Li, K., et al. (2012). Global landscape of HIV–human protein complexes. Nature 481, 365–370. 10.1038/nature10719.

31. Steinkamp, R., Tsitsiridis, G., Brauner, B., Montrone, C., Fobo, G., Frishman, G., Avram, S., Oprea, T.I., and Ruepp, A. (2024). CORUM in 2024: protein complexes as drug targets. Nucleic Acids Res. 53, D651–D657. 10.1093/nar/gkae1033.

32. Aeberhard, L., Banhart, S., Fischer, M., Jehmlich, N., Rose, L., Koch, S., Laue, M., Renard, B.Y., Schmidt, F., and Heuer, D. (2015). The Proteome of the Isolated Chlamydia trachomatis Containing Vacuole Reveals a Complex Trafficking Platform Enriched for Retromer Components. PLoS Pathog. 11, e1004883. 10.1371/journal.ppat.1004883.

33. Olson, M.G., Widner, R.E., Jorgenson, L.M., Lawrence, A., Lagundzin, D., Woods, N.T., Ouellette, S.P., and Rucks, E.A. (2019). Proximity Labeling To Map Host-Pathogen Interactions at the Membrane of a Bacterium-Containing Vacuole in Chlamydia trachomatis-Infected Human Cells. Infect. Immun. 87, e00537–19. 10.1128/iai.00537-19.

34. Dickinson, M.S., Anderson, L.N., Webb-Robertson, B.-J.M., Hansen, J.R., Smith, R.D., Wright, A.T., and Hybiske, K. (2019). Proximity-dependent proteomics of the Chlamydia trachomatis inclusion membrane reveals functional interactions with endoplasmic reticulum exit sites. PLoS Pathog. 15, e1007698. 10.1371/journal.ppat.1007698.

35. Shannon, P., Markiel, A., Ozier, O., Baliga, N.S., Wang, J.T., Ramage, D., Amin, N., Schwikowski, B., and Ideker, T. (2003). Cytoscape: A Software Environment for Integrated Models of Biomolecular Interaction Networks. Genome Res. 13, 2498–2504. 10.1101/gr.1239303.

36. Weber, M.M., Noriea, N.F., Bauler, L.D., Lam, J.L., Sager, J., Wesolowski, J., Paumet, F., and Hackstadt, T. (2016). A Functional Core of IncA Is Required for Chlamydia trachomatis Inclusion Fusion. J Bacteriol 198, 1347–1355. 10.1128/jb.00933-15.

37. Faris, R., Merling, M., Andersen, S.E., Dooley, C.A., Hackstadt, T., and Weber, M.M. (2019). Chlamydia trachomatis CT229 Subverts Rab GTPase-Dependent CCV Trafficking Pathways to Promote Chlamydial Infection. Cell Reports 26, 3380–3390.e5. 10.1016/j.celrep.2019.02.079.

38. Chellas Géry, B., Linton, C.N., and Fields, K.A. (2007). Human GCIP interacts with CT847, a novel Chlamydia trachomatis type III secretion substrate, and is degraded in a tissue culture infection model. Cell Microbiol 9, 2417–2430. 10.1111/j.1462-5822.2007.00970.x.

39. Pennini, M.E., Perrinet, S., Dautry-Varsat, A., and Subtil, A. (2010). Histone Methylation by NUE, a Novel Nuclear Effector of the Intracellular Pathogen Chlamydia trachomatis. Plos Pathog 6, e1000995. 10.1371/journal.ppat.1000995.

40. Hamaoui, D., Cossé, M.M., Mohan, J., Lystad, A.H., Wollert, T., and Subtil, A. (2020). The Chlamydia effector CT622/TaiP targets a nonautophagy related function of ATG16L1. Proc. Natl. Acad. Sci. 117, 26784–26794. 10.1073/pnas.2005389117.

41. Dolat, L., Carpenter, V.K., Chen, Y.-S., Suzuki, M., Smith, E.P., Kuddar, O., and Valdivia, R.H. (2022). Chlamydia repurposes the actin-binding protein EPS8 to disassemble epithelial tight junctions and promote infection. Cell Host Microbe 30, 1685–1700.e10. 10.1016/j.chom.2022.10.013.

42. Fu, M., Liu, Y., Wang, G., Wang, P., Zhang, J., Chen, C., Zhao, M., Zhang, S., Jiao, J., Ouyang, X., et al. (2022). A protein–protein interaction map reveals that the Coxiella burnetii effector CirB inhibits host proteasome activity. PLoS Pathog. 18, e1010660. 10.1371/journal.ppat.1010660.

43. Walch, P., Selkrig, J., Knodler, L.A., Rettel, M., Stein, F., Fernandez, K., Viéitez, C., Potel, C.M., Scholzen, K., Geyer, M., et al. (2021). Global mapping of Salmonella enterica-host protein-protein interactions during infection. Cell Host Microbe 29, 1316–1332.e12. 10.1016/j.chom.2021.06.004.

44. Penn, B.H., Netter, Z., Johnson, J.R., Dollen, J.V., Jang, G.M., Johnson, T., Ohol, Y.M., Maher, C., Bell, S.L., Geiger, K., et al. (2018). An Mtb-Human Protein-Protein Interaction Map Identifies a Switch between Host Antiviral and Antibacterial Responses. Mol. Cell 71, 637–648.e5. 10.1016/j.molcel.2018.07.010.

45. Hower, S., Wolf, K., and Fields, K.A. (2009). Evidence that CT694 is a novel Chlamydia trachomatis T3S substrate capable of functioning during invasion or early cycle development. Mol. Microbiol. 72, 1423–1437. 10.1111/j.1365-2958.2009.06732.x.

46. Ooij, C. van, Apodaca, G., and Engel, J. (1997). Characterization of the Chlamydia trachomatis vacuole and its interaction with the host endocytic pathway in HeLa cells. Infect. Immun. 65, 758–766. 10.1128/iai.65.2.758-766.1997.

47. Rzomp, K.A., Scholtes, L.D., Briggs, B.J., Whittaker, G.R., and Scidmore, M.A. (2003). Rab GTPases Are Recruited to Chlamydial Inclusions in Both a Species-Dependent and Species- Independent Manner. Infect Immun 71, 5855–5870. 10.1128/iai.71.10.5855.

48. Beck, M., and Hurt, E. (2017). The nuclear pore complex: understanding its function through structural insight. Nat. Rev. Mol. Cell Biol. 18, 73–89. 10.1038/nrm.2016.147.

49. Pritchard, C.E.J., Fornerod, M., Kasper, L.H., and Deursen, J.M.A. van (1999). RAE1 Is a Shuttling mRNA Export Factor That Binds to a GLEBS-like NUP98 Motif at the Nuclear Pore Complex through Multiple Domains. J. Cell Biol. 145, 237–254. 10.1083/jcb.145.2.237.

50. Schott, B.H., Antonia, A.L., Wang, L., Pittman, K.J., Sixt, B.S., Barnes, A.B., Valdivia, R.H., and Ko, D.C. (2020). Modeling of variables in cellular infection reveals CXCL10 levels are regulated by human genetic variation and the Chlamydia-encoded CPAF protease. Sci Rep- uk 10, 18269. 10.1038/s41598-020-75129-y.

51. Ouellette, S.P. (2018). Feasibility of a Conditional Knockout System for Chlamydia Based on CRISPR Interference. Front. Cell. Infect. Microbiol. 8, 59. 10.3389/fcimb.2018.00059.

52. Gong, D., Kim, Y.H., Xiao, Y., Du, Y., Xie, Y., Lee, K.K., Feng, J., Farhat, N., Zhao, D., Shu, S., et al. (2016). A Herpesvirus Protein Selectively Inhibits Cellular mRNA Nuclear Export. Cell Host Microbe 20, 642–653. 10.1016/j.chom.2016.10.004.

53. Shen, Q., Kumari, S., Xu, C., Jang, S., Shi, J., Burdick, R.C., Levintov, L., Xiong, Q., Wu, C., Devarkar, S.C., et al. (2023). The capsid lattice engages a bipartite NUP153 motif to mediate nuclear entry of HIV-1 cores. Proc. Natl. Acad. Sci. 120, e2202815120. 10.1073/pnas.2202815120.

54. Dickson, C.F., Hertel, S., Tuckwell, A.J., Li, N., Ruan, J., Al-Izzi, S.C., Ariotti, N., Sierecki, E., Gambin, Y., Morris, R.G., et al. (2024). The HIV capsid mimics karyopherin engagement of FG-nucleoporins. Nature 626, 836–842. 10.1038/s41586-023-06969-7.

55. Quan, B., Seo, H.-S., Blobel, G., and Ren, Y. (2014). Vesiculoviral matrix (M) protein occupies nucleic acid binding site at nucleoporin pair (Rae1•Nup98). Proc. Natl. Acad. Sci. 111, 9127–9132. 10.1073/pnas.1409076111.

56. Faria, P.A., Chakraborty, P., Levay, A., Barber, G.N., Ezelle, H.J., Enninga, J., Arana, C., Deursen, J. van, and Fontoura, B.M.A. (2005). VSV Disrupts the Rae1/mrnp41 mRNA Nuclear Export Pathway. Mol. Cell 17, 93–102. 10.1016/j.molcel.2004.11.023.

57. Miorin, L., Kehrer, T., Sanchez-Aparicio, M.T., Zhang, K., Cohen, P., Patel, R.S., Cupic, A., Makio, T., Mei, M., Moreno, E., et al. (2020). SARS-CoV-2 Orf6 hijacks Nup98 to block STAT nuclear import and antagonize interferon signaling. Proc. Natl. Acad. Sci. 117, 28344–28354. 10.1073/pnas.2016650117.

58. Monette, A., Panté, N., and Mouland, A.J. (2011). HIV-1 remodels the nuclear pore complex. J. Cell Biol. 193, 619–631. 10.1083/jcb.201008064.

59. Ibana, J.A., Sherchand, S.P., Fontanilla, F.L., Nagamatsu, T., Schust, D.J., Quayle, A.J., and Aiyar, A. (2018). Chlamydia trachomatis-infected cells and uninfected-bystander cells exhibit diametrically opposed responses to interferon gamma. Sci. Rep. 8, 8476. 10.1038/s41598-018-26765-y.

60. Nunzio, F.D., Fricke, T., Miccio, A., Valle-Casuso, J.C., Perez, P., Souque, P., Rizzi, E., Severgnini, M., Mavilio, F., Charneau, P., et al. (2013). Nup153 and Nup98 bind the HIV-1 core and contribute to the early steps of HIV-1 replication. Virology 440, 8–18. 10.1016/j.virol.2013.02.008.

61. 61. Lad, S.P., Fukuda, E.Y., Li, J., Maza, L.M. de la, and Li, E. (2005). Up-Regulation of the JAK/STAT1 Signal Pathway during Chlamydia trachomatis Infection. J. Immunol. 174, 7186– 7193. 10.4049/jimmunol.174.11.7186.

62. Fontanilla, F.L., Ibana, J.A., Carabeo, R.A., and Brinkworth, A.J. (2024). Chlamydia trachomatis modulates the expression of JAK-STAT signaling components to attenuate the type II interferon response of epithelial cells. mBio 15, e01834–24. 10.1128/mbio.01834-24.

63. Haldar, A.K., Piro, A.S., Finethy, R., Espenschied, S.T., Brown, H.E., Giebel, A.M., Frickel, E.M., Nelson, D.E., and Coers, J. (2016). Chlamydia trachomatis Is Resistant to Inclusion Ubiquitination and Associated Host Defense in Gamma Interferon-Primed Human Epithelial Cells. mBio 7, e01417–16. 10.1128/mbio.01417-16.

64. Giebel, A.M., Hu, S., Rajaram, K., Finethy, R., Toh, E., 1, J.A.B., Morrison, S.G., Suchland, R.J., Stein, B.D., Coers, J., et al. (2019). Genetic Screen in Chlamydia muridarum Reveals Role for an Interferon-Induced Host Cell Death Program in Antimicrobial Inclusion Rupture. mBio 10, ]e00385-19. 10.1128/mbio.00385-19.

65. Walsh, S.C., Reitano, J.R., Dickinson, M.S., Kutsch, M., Hernandez, D., Barnes, A.B., Schott, B.H., Wang, L., Ko, D.C., Kim, S.Y., et al. (2022). The bacterial effector GarD shields Chlamydia trachomatis inclusions from RNF213-mediated ubiquitylation and destruction. Cell Host Microbe 30, 1671–1684.e9. 10.1016/j.chom.2022.08.008.

66. Morrison, R.P. (2000). Differential sensitivities of Chlamydia trachomatis strains to inhibitory effects of gamma interferon. Infect Immun 68, 6038–6040. 10.1128/iai.68.10.6038-6040.2000.

67. Fontanilla, F.L., Carabeo, R.A., and Brinkworth, A.J. (2024). Chlamydia trachomatis modulates the expression of JAK-STAT signaling components to attenuate the Type II interferon response of epithelial cells. bioRxiv, 2024.01.09.574898. 10.1101/2024.01.09.574898.

68. Barta, M.L., Hickey, J., Kemege, K.E., Lovell, S., Battaile, K.P., and Hefty, P.S. (2013). Structure of CT584 from Chlamydia trachomatis refined to 3.05 Å resolution. Acta Crystallogr. Sect. F: Struct. Biol. Cryst. Commun. 69, 1196–1201. 10.1107/s1744309113027371.

69. Pha, K., Mirrashidi, K., Sherry, J., Tran, C.J., Herrera, C.M., McMahon, E., Elwell, C.A., and Engel, J.N. (2024). The Chlamydia effector IncE employs two short linear motifs to reprogram host vesicle trafficking. Cell Rep. 43, 114624. 10.1016/j.celrep.2024.114624.

70. Pittner, N.A., Solomon, R.N., Bui, D.-C., and McBride, J.W. (2023). Ehrlichia effector SLiM-icry: Artifice of cellular subversion. Front. Cell. Infect. Microbiol. 13, 1150758. 10.3389/fcimb.2023.1150758.

71. Mojica, S.A., Hovis, K.M., Frieman, M.B., Tran, B., Hsia, R., Ravel, J., Jenkins-Houk, C., Wilson, K.L., and Bavoil, P.M. (2015). SINC, a type III secreted protein of Chlamydia psittaci, targets the inner nuclear membrane of infected cells and uninfected neighbors. Mol Biol Cell 26, 1918–1934. 10.1091/mbc.e14-11-1530.

72. Fleming, A., Sampey, G., Chung, M., Bailey, C., Hoek, M.L., Kashanchi, F., and Hakami, R.M. (2014). The carrying pigeons of the cell: exosomes and their role in infectious diseases caused by human pathogens. Pathog. Dis. 71, 109–120. 10.1111/2049-632x.12135.

73. Jahnke, R., Matthiesen, S., Zaeck, L.M., Finke, S., and Knittler, M.R. (2022). Chlamydia trachomatis Cell-to-Cell Spread through Tunneling Nanotubes. Microbiol. Spectr. 10, e02817–22. 10.1128/spectrum.02817-22.

74. Grieshaber, S.S., Grieshaber, N.A., Miller, N., and Hackstadt, T. (2006). Chlamydia trachomatis Causes Centrosomal Defects Resulting in Chromosomal Segregation Abnormalities. Traffic 7, 940–949. 10.1111/j.1600-0854.2006.00439.x.

75. Scanlon, K.R., Keb, G., Wolf, K., Jewett, T.J., and Fields, K.A. (2023). Chlamydia trachomatis TmeB antagonizes actin polymerization via direct interference with Arp2/3 activity. Front. Cell. Infect. Microbiol. 13, 1232391. 10.3389/fcimb.2023.1232391.

76. Chhabra, E.S., and Higgs, H.N. (2006). INF2 Is a WASP Homology 2 Motif-containing Formin That Severs Actin Filaments and Accelerates Both Polymerization and Depolymerization*. J. Biol. Chem. 281, 26754–26767. 10.1074/jbc.m604666200.

77. Truong, D., Copeland, J.W., and Brumell, J.H. (2014). Bacterial subversion of host cytoskeletal machinery: Hijacking formins and the Arp2/3 complex. BioEssays 36, 687–696. 10.1002/bies.201400038.

