## Supplementary material for "The *Chlamydia trachomatis* secreted effector CebN targets nucleoporins and Rae1 to antagonize STAT1 nuclear import": S1 Table Primers

Table S1: Primers used in this study. Nucleotides in bold correspond to the restriction site, and nucleotides in blue correspond to the FLAG-tag. IP = constructs used for immunoprecipitation, IF = constructs used for immunofluorescence

| **Primer Name** | **Sequence** | **Uses** |
| --- | --- | --- |
| **Expression in *C.t.*** | | |
| CT016 NotI F | CC**GCGGCCGC**ATGAAAGTCAAAATTAATGATCAGTTC | AP-MS |
| CT016 FLAG KpnI R | CC**GGTACCTTACTTATCGTCGTCATCCTTGTAATC**AGTATAAAGAACAGCTTTCACGTGTTC | AP-MS |
| CT053 NotI F | CC**GCGGCCGC**ATGAAAAGTGAGCGTTTAAAAAAATT | AP-MS |
| CT053 FLAG KpnI R | CC**GGTACCTTACTTATCGTCGTCATCCTTGTAATC**CCATTCATTCGCGTCAGG | AP-MS |
| CT142 NotI F | CC**GCGGCCGC**ATGAGTGATTCTGACAAAATTATTAAT | AP-MS |
| CT142 FLAG KpnI R | CC**GGTACCTTACTTATCGTCGTCATCCTTGTAATC**TCCTCCTATCTCTGGGTATACGAG | AP-MS |
| CT143 NotI F | CC**GCGGCCGC**ATGAAGAAACCAGTATTTACAGGGG | AP-MS |
| CT143 FLAG KpnI R | CC**GGTACCTTACTTATCGTCGTCATCCTTGTAATC**ATCTGCCTCCTTATAAGAAGAACCA | AP-MS |
| CT144 NotI F | CC**GCGGCCGC**ATGACAACACCAGATAATAATACTATTGAT | AP-MS |
| CT144 FLAG KpnI R | CC**GGTACCTTACTTATCGTCGTCATCCTTGTAATC**AGGAACAACAGGTAGCCGAA | AP-MS |
| CT161 NotI F | CC**GCGGCCGC**GTGGCTAGAAAACCTTTAGTAGATAGA | AP-MS |
| CT161 FLAG KpnI R | CC**GGTACCTTACTTATCGTCGTCATCCTTGTAATC**GTCATAAAAATTTTCCATTTCTGTAGG | AP-MS |
| CT311 NotI F | CC**GCGGCCGC**ATGAAAAGAGTTATCCTCTGCTCTCT | AP-MS |
| CT311 FLAG KpnI R | CC**GGTACCTTACTTATCGTCGTCATCCTTGTAATC**TTTTCCATTTTGCAGATCTTTCA | AP-MS |
| CT386 NotI F | CC**GCGGCCGC**ATGCAAATTCCAAGAAGTGTTG | AP-MS |
| CT386 FLAG KpnI R | CC**GGTACCTTACTTATCGTCGTCATCCTTGTAATC**TACTAATCTCTGCTGTTTTAACA | AP-MS |
| CT392 NotI F | CC**GCGGCCGC**ATGTCTTCTATACAAGGAAC | AP-MS |
| CT392 FLAG KpnI R | CC**GGTACCTTACTTATCGTCGTCATCCTTGTAATC**AAATCCTCTATCATCATCGG | AP-MS |
| CT504 NotI F | CC**GCGGCCGC**GTGTATTTTACAAGAGATCCAGTCAT | AP-MS |
| CT504 FLAG KpnI R | CC**GGTACCTTACTTATCGTCGTCATCCTTGTAATC**CTCTTCTGAAGAAATACTGTC | AP-MS |
| CT537 NotI F | CC**GCGGCCGC**ATGGGTAGATACAGAAGAG | AP-MS |
| CT537 FLAG KpnI R | CC**GGTACC**ATCGTTCTCCATAAAAAAGC | AP-MS |
| CT583 NotI F | CC**GCGGCCGC**ATGGGAAATATTAAAACCCTTTTAGAG | AP-MS |
| CT583 FLAG KpnI R | CC**GGTACCTTACTTATCGTCGTCATCCTTGTAATC**TCGATTTCTAGAGTTTTGGGTTT | AP-MS |
| CT584 (CebN) NotI F | CC**GCGGCCGC**ATGACGACGAAACCCAAAACTCT | AP-MS, IP, IF |
| CT584 (CebN) FLAG KpnI R | CC**GGTACCTTACTTATCGTCGTCATCCTTGTAATC**CACAGATTTCGTTAATTCTTCAA | AP-MS, IP, IF |
| CT584 (CebN) 1-160aa FLAG KpnI R | CC**GGTACCTTACTTATCGTCGTCATCCTTGTAATC**TTTTTCTCCGATAGGATTTTT | IP, IF |
| CT584 (CebN) 1-140aa FLAG KpnI R | CC**GGTACCTTACTTATCGTCGTCATCCTTGTAATC**ATGGCAGGTATTTTGTACATCCG | IP, IF |
| CT584 (CebN) 1-100aa FLAG KpnI R | CC**GGTACCTTACTTATCGTCGTCATCCTTGTAATC**GAATTCACCGCGCTCATGG | IP, IF |
| CT606.1 NotI F | CC**GCGGCCGC**TTGGAAGATAGAATGATCGACGG | AP-MS |
| CT606.1 FLAG KpnI R | CC**GGTACCTTACTTATCGTCGTCATCCTTGTAATC**CTCGCGGGGAAAGAGAGTCT | AP-MS |
| CT610 NotI F | CC**GCGGCCGC**ATGATGGAGGTGTTTATGAAT | AP-MS |
| CT610 FLAG KpnI R | CC**GGTACCTTACTTATCGTCGTCATCCTTGTAATC**ATAAGATTGATGACAACTACAAC | AP-MS |
| CT620 NotI F | CC**GCGGCCGC**ATGTGTTCTATGAACATATTTAATAAAATTAACTC | AP-MS |
| CT620 FLAG SalI R | CC**GTCGACTTACTTATCGTCGTCATCCTTGTAATC**ACTAGCCAGTTTTCTTGTTAAACCA | AP-MS |
| CT621 NotI F | CC**GCGGCCGC**ATGAACCGTATTCATCGTACACAA | AP-MS |
| CT621 FLAG SalI R | CC**GTCGACTTACTTATCGTCGTCATCCTTGTAATC**TCTTAAGAGATTACGCGCTAATCC | AP-MS |
| CT622 NotI F | CC**GCGGCCGC**ATGGAATCAGGACCAGAATCAG | AP-MS |
| CT622 FLAG SalI R | CC**GTCGACTTACTTATCGTCGTCATCCTTGTAATC**AGAAAGATAACCAGAGAATAGAGAA | AP-MS |
| CT627 NotI F | CC**GCGGCCGC**ATGGAAAAGAATTATTATGC | AP-MS |
| CT627 FLAG KpnI R | CC**GGTACCTTACTTATCGTCGTCATCCTTGTAATC**TGCTGGTTGGTTTTCTTGTTCTT | AP-MS |
| CT631 NotI F | CC**GCGGCCGC**ATGAAAACGTTAATTGATAACA | AP-MS |
| CT631 FLAG KpnI R | CC**GGTACCTTACTTATCGTCGTCATCCTTGTAATC**TAAACAAATAATTCCTTCAAACT | AP-MS |
| CT652.1 NotI F | CC**GCGGCCGC**ATGGACCAGTTATCACAGA | AP-MS |
| CT652.1 FLAG KpnI R | CC**GGTACCTTACTTATCGTCGTCATCCTTGTAATC**ACCTTGGGAATCTTCTT | AP-MS |
| CT656 NotI F | CC**GCGGCCGC**ATGGACACGCAATTCATAGCG | AP-MS |
| CT656 FLAG KpnI R | CC**GGTACCTTACTTATCGTCGTCATCCTTGTAATC**ATCTCTGTATACCGAACGCATTT | AP-MS |
| CT671 NotI F | CC**GCGGCCGC**ATGGAATTAAATAAAACTTCGGAATCT | AP-MS |
| CT671 FLAG KpnI R | CC**GTCGACTTACTTATCGTCGTCATCCTTGTAATC**TATATGAGCTTCTTCTACTTTCTTCTC | AP-MS |
| CT691 NotI F | CC**GCGGCCGC**ATGCAAGTCCTAGCAAGTTTATT | AP-MS |
| CT691 FLAG KpnI R | CC**GTCGACTTACTTATCGTCGTCATCCTTGTAATC**TTTCTCTTCTAGCGTCATACTCA | AP-MS |
| CT694 (TmeA) NotI F | CC**GCGGCCGC**ATGAGTATTCGACCTACTAATGGGAG | IP, IF |
| CT694 (TmeA) FLAG SalI | CC**GTCGACTTACTTATCGTCGTCATCCTTGTAATC**GTCTAAGAAAACAGAAGAAGTTATGAC | IP, IF |
| CT695 NotI F | CC**GCGGCCGC**GTGAGTAGCATAAGCCCTATAGGG | AP-MS |
| CT695 FLAG KpnI R | CC**GTCGACTTACTTATCGTCGTCATCCTTGTAATC**GATATTCCCAACCGAAGAAGG | AP-MS |
| CT711 NotI F | CC**GCGGCCGC**GTGTCAATACAACCTACATCCATTTC | AP-MS |
| CT711 FLAG KpnI R | CC**GGTACCTTACTTATCGTCGTCATCCTTGTAATC**TTTAAATCTACGGATCAACTTAGCAA | AP-MS |
| CT712 NotI F | CC**GCGGCCGC**ATGAGAAACCATCCGATTCC | AP-MS |
| CT712 FLAG KpnI R | CC**GGTACCTTACTTATCGTCGTCATCCTTGTAATC**GCTAGAAGCCAATGTTCTATATACATT | AP-MS |
| CT736 NotI F | CC**GCGGCCGC**ATGCAACTCACCTCACAAG | AP-MS |
| CT736 FLAG KpnI R | CC**GTCGACTTACTTATCGTCGTCATCCTTGTAATC**GTCTTTTTCATATGTGCCC | AP-MS |
| CT738 NotI F | CC**GCGGCCGC**ATGGAAGGTTTTTTCCCTATA | AP-MS |
| CT738 FLAG SalI R | CC**GGTACCTTACTTATCGTCGTCATCCTTGTAATC**TACAAGGCTCGGAGCAGAAAAGA | AP-MS |
| CT848 NotI F | CC**GCGGCCGC**ATGTGGCATAAAGAACCAATGTATG | AP-MS |
| CT848 FLAG SalI R | CC**GTCGACTTACTTATCGTCGTCATCCTTGTAATC**GATATTCGCGATCAAGCTAACG | AP-MS |
| CT849 NotI F | CC**GCGGCCGC**ATGTCAGCACCAACCTCACA | AP-MS |
| CT849 FLAG SalI R | CC**GTCGACTTACTTATCGTCGTCATCCTTGTAATC**AGACAGGGGTTTATTTAATTGGTTAAC | AP-MS |
|  | **Ectopic Expression** |  |
| CT584 (CebN) KpnI F | CC**GGTACC**ATGACGACGAAACCCAAAACTCT | IF |
| CT584 (CebN) SalI R | CC**GTCGAC**TTACACAGATTTCGTTAATTCTT | IF |
| CT694 (TmeA) SalI F | CC**GGTACC**ATGAGTATTCGACCTACTAATGGGAG | IF |
| CT694 (TmeA) XhoI R | CC**CTCGAGTTA**GTCTAAGAAAACAGAAGAAGTT | IF |
| **Mutagenesis** | | |
| CebN 186 CRISPRi | tgaatataattttaattatatcacgcactagctcagattcagtagaccgctgttgATAAAACGAAAGGCCCAGTCTTTCGACTGAGCCTTTCGTTTTATTTG**GGCGTGGCAAAAATTCTTTT**ATCTACAAGAGTAGAAATTAGGTGTCATTATAAGAAAACCGTCTCCACCGGTGAAtgcatgatctacgtgcgtcacatgcagtaccgacgtcttaagacccactttcaca | CRISPRi |
| CepN KD sRNA loop | 5' ttagtgttagaaagaaggttagttgtaact 3' | sRNA-mediated KD |
