## Supplementary material for "The *Chlamydia trachomatis* secreted effector CebN targets nucleoporins and Rae1 to antagonize STAT1 nuclear import": Supp Figures

**SUPPLEMENTAL MATERIALS**

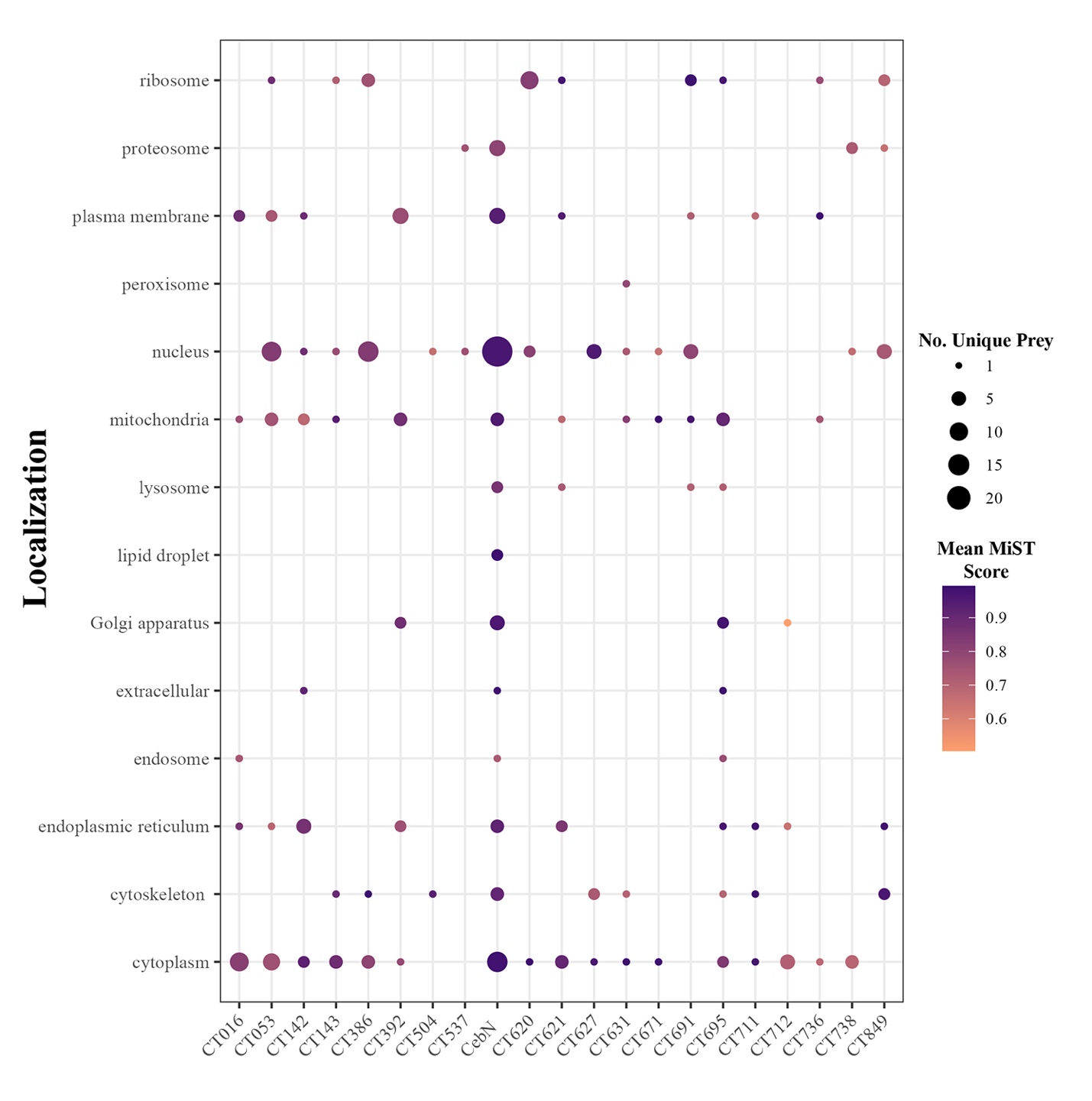

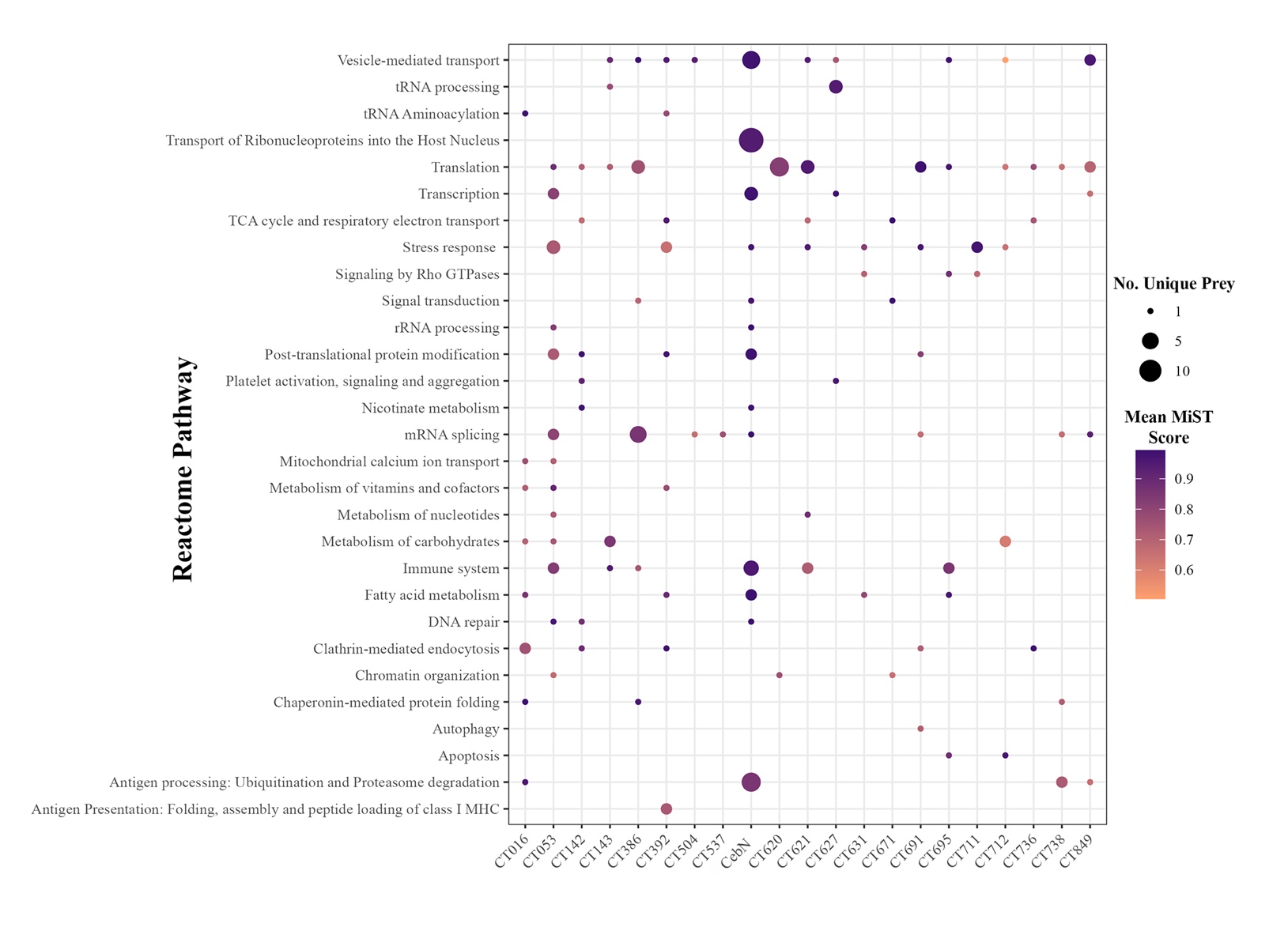
 **Fig. S1.** Subcellular localization and Reactome pathways targeted by *C.t.* secreted effector proteins. MiST was used to identify high-confidence interacting partners for each effector screened. Proteins with MiST score ≥0.7 were considered significant and analyzed for (A) subcellular localization and (B) Reactome pathways using UniProt and Genecards. Dot size corresponds to the number of unique prey(s) and color indicates the average MiST score of those prey.

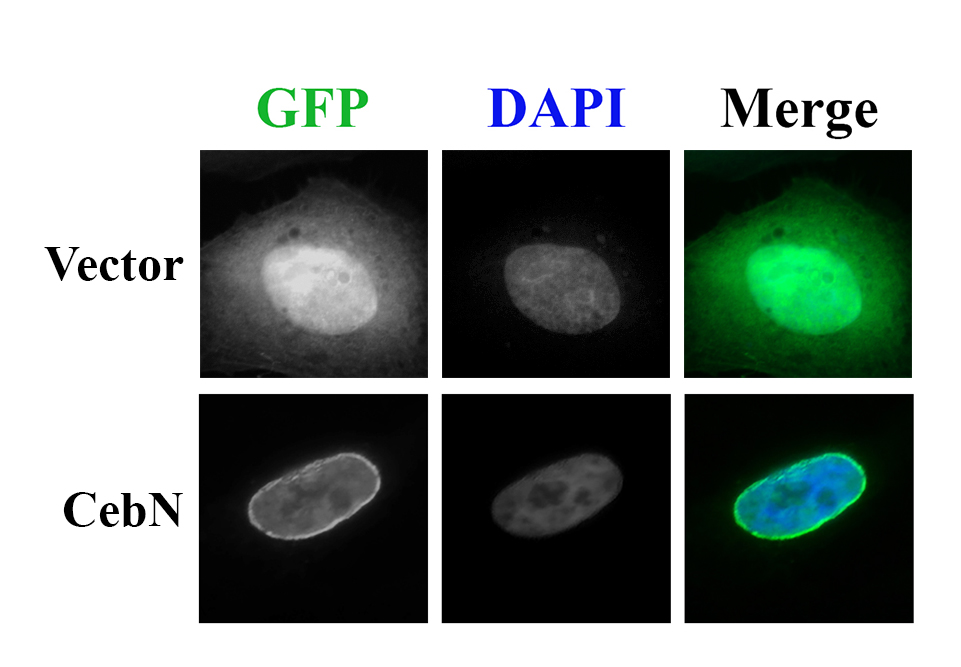

**Fig. S2**. CebN localizes to the nuclear envelope. HeLa cells were transfected with GFP-CebN or GFP (green), fixed, and stained with DAPI (blue) to demark to the nucleus. Data are representative of three independent experiments.

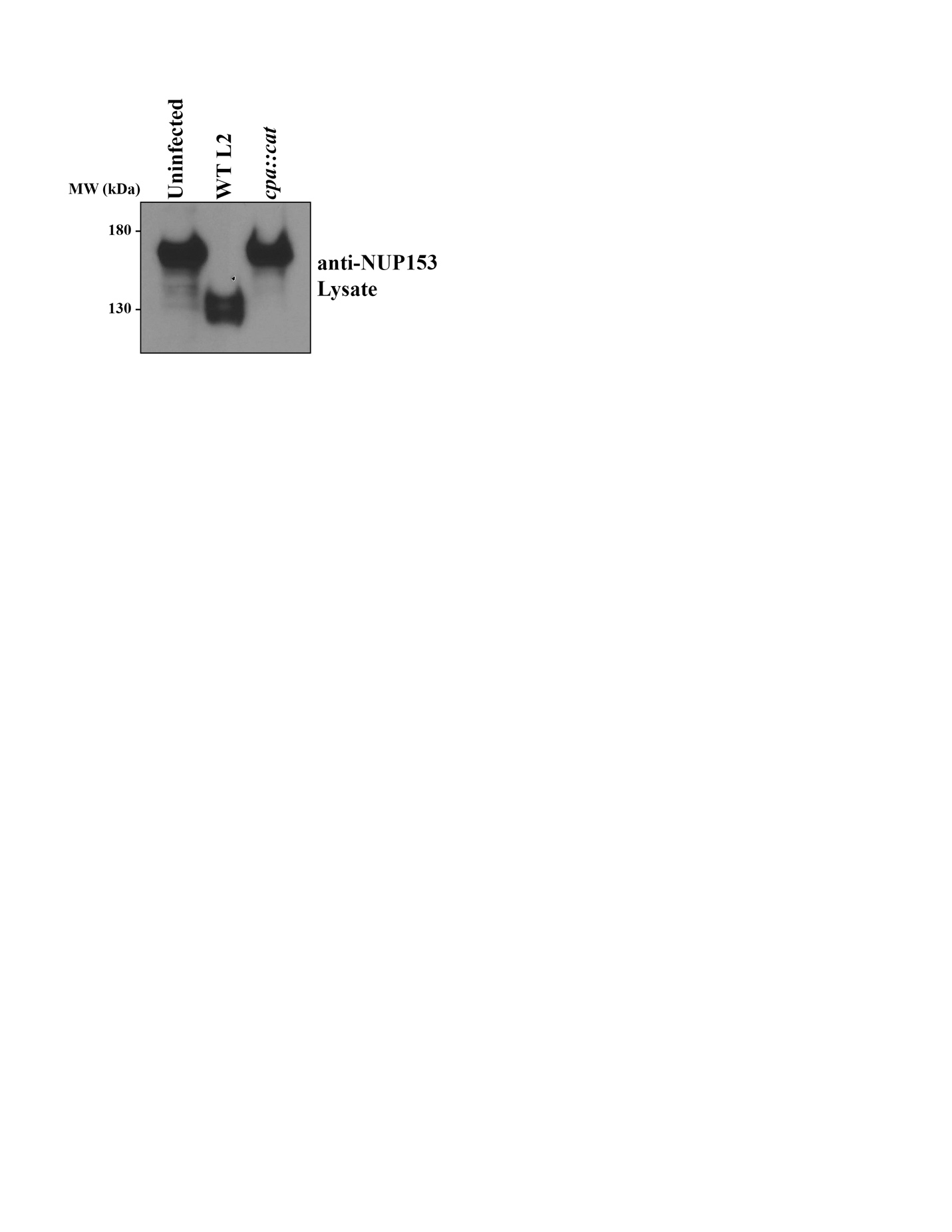

**Fig. S3.** Proteolytic cleavage of NUP153 is an artifact of CPAF activity during sample processing. HeLa cells were left uninfected or infected with WT L2 or a CPAF mutant (*cpa::cat*). Cell lysates were analyzed by immunoblotting with antibodies against NUP153.

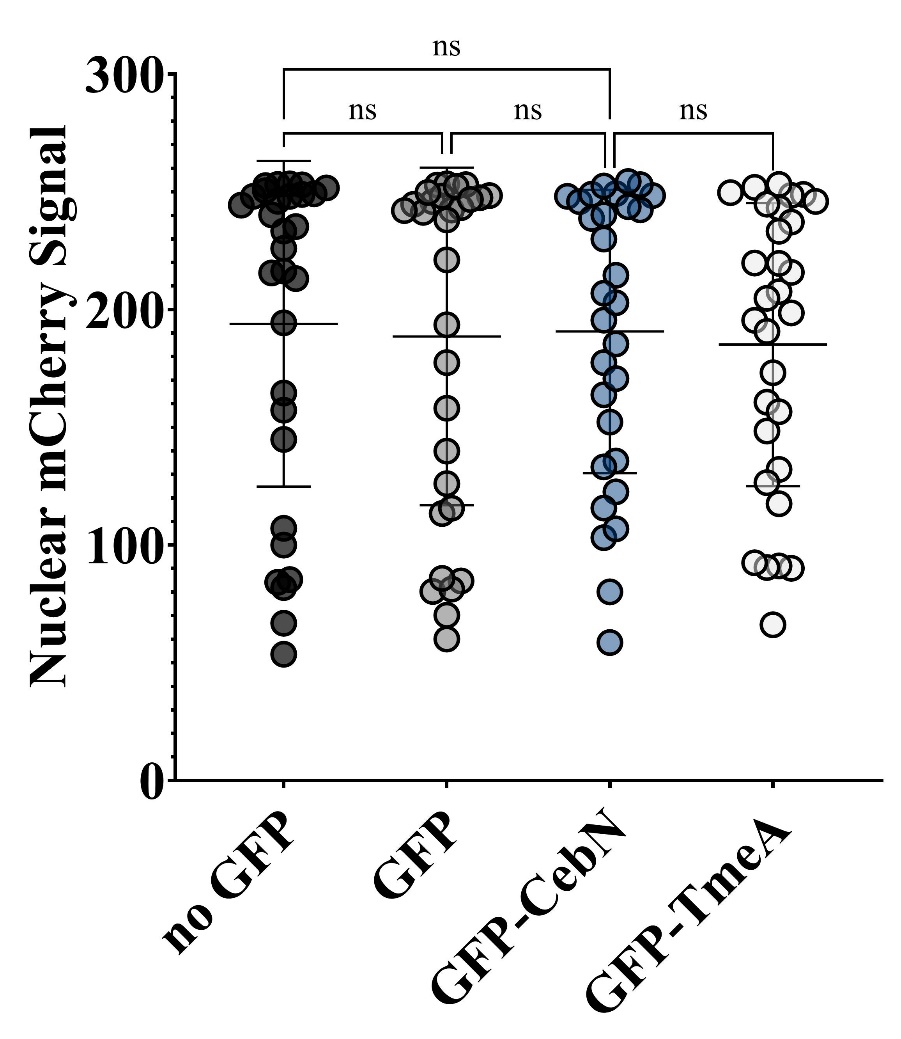

**Fig. S4.** CebN does not inhibit nuclear translocation of NLS-tagged mCherry. HeLa cells were transfected with mCherry-NLS alone or co-transfected with mCherry-NLS and GFP empty vector, GFP-CebN, or GFP-TmeA. Cells were fixed and stained with DAPI. Nuclear mCherry signal intensity was quantified and plotted as relative fluorescence units (RFU). Error bars represent standard deviation. Statistical significance was determined using one-way ANOVA followed by Tukey’s multiple comparisons test. Data are representative of three replicates.
