## Supplementary material for "The *Chlamydia trachomatis* secreted effector CebN targets nucleoporins and Rae1 to antagonize STAT1 nuclear import": S2 Antibodies

Table S2: Primary and secondary antibodies used in this study. WB = western blot, IF = immunofluorescence.

| **Target** | **Company** | **Product No.** | **Use** | **Dilution** |
| --- | --- | --- | --- | --- |
| CyaA | Santa Cruz Biotech | SC-13582 | Western blotting | 1:500 |
| BlaM (β lactamase) | QED | 15720 | Western blotting | 1:2000 |
| GSK-3β | Cell Signaling | 9325S | Western blotting | 1:2000 |
| Phospho-GSK-3β | Cell Signaling | 9336S | Western blotting | 1:2000 |
| FLAG | Invitrogen | 701629 | Western blotting | 1:5000 |
| GFP | Novus | NB600-597 | Immunofluorescence | 1:1000 |
| NUP54 | Proteintech | 16232-1-AP | Western blotting, Immunofluorescence | WB 1:2000  IF 1:1000 |
| NUP153 | Novus | NBP1-81725 | Western blotting, Immunofluorescence | WB 1:2000  IF 1:1000 |
| NUP214 | Abcam | AB70497 | Western blotting, Immunofluorescence | WB 1:2000  IF 1:1000 |
| Rae1 | Novus | NBP1-31027 | Western blotting | 1:2000 |
| FLAG | Cell Signaling | 14793T | Immunofluorescence | 1:1000 |
| *C.t.* HSP60 | Sigma | MABF2108 | Immunofluorescence | 1:1000 |
| STAT1 | Cell Signaling | 14994T | Immunofluorescence | 1:1000 |
| Goat anti-Mouse IgG (H+L) Secondary Antibody, HRP | Bio-Rad | 1706516 | Western blotting | 1:10,000 |
| Goat Anti-Rabbit IgG (H + L)-HRP Conjugate | Bio-Rad | 1706515 | Western blotting | 1:10,000 |
| Goat anti-Mouse IgG (H+L) Cross-Adsorbed Secondary Antibody, Alexa Fluor™ 488 | Invitrogen | A11001 | Immunofluorescence | 1:1000 |
| Abberior STAR 635 | Abberior | ST635-1002 | Immunofluorescence | 1:250 |
| Goat anti-Mouse IgG (H+L) Cross-Adsorbed Secondary Antibody, DyLight™ 594 | Invitrogen | 35511 | Immunofluorescence | STED 1:250  IF 1:1000 |
| Goat anti-Rabbit IgG (H+L) Highly Cross-Adsorbed Secondary Antibody, Alexa Fluor™ Plus 647 | Invitrogen | A32733 | Immunofluorescence | 1:1000 |
| Goat anti-Rabbit IgG (H+L) Cross-Adsorbed Secondary Antibody, DyLight™ 594 | Invitrogen | 35561 | Immunofluorescence | 1:1000 |
